# Rolosense: Mechanical detection of SARS-CoV-2 using a DNA-based motor

**DOI:** 10.1101/2023.02.27.530294

**Authors:** Selma Piranej, Luona Zhang, Alisina Bazrafshan, Mariana Marin, Gregory B. Melikyan, Khalid Salaita

## Abstract

Assays detecting viral infections play a significant role in limiting the spread of diseases such as SARS-CoV-2. Here we present Rolosense, a virus sensing platform that transduces the motion of synthetic DNA-based motors transporting 5-micron particles on RNA fuel chips. Motors and chips are modified with virus-binding aptamers that lead to stalling of motion. Therefore, motors perform a “mechanical test” of viral target and stall in the presence of whole virions which represents a unique mechanism of transduction distinct from conventional assays. Rolosense can detect SARS-CoV-2 spiked in artificial saliva and exhaled breath condensate with a sensitivity of 10^3^ copies/mL and discriminates among other respiratory viruses. The assay is modular and amenable to multiplexing, as we demonstrated one-pot detection of influenza A and SARS-CoV-2. As a proof-of-concept, we show readout can be achieved using a smartphone camera in as little as 15 mins without any sample preparation steps. Taken together, mechanical detection using Rolosense can be broadly applied to any viral target and has the potential to enable rapid, low-cost, point-of-care screening of circulating viruses.

## Main

Virus sensing is primarily performed using nucleic acid-based assays or alternatively by detecting protein antigens using colorimetric, fluorogenic, or electrochemical reporters. The gold-standard diagnostic for SARS-CoV-2 infection is RT-qPCR which detects down to ∼10^2-3^ viral RNA copies/mL and is typically performed at central facilities with a 10–15-hour turnaround time.^1,2^ Alternate nucleic acid-based diagnostics include isothermal amplification such as loop-mediated isothermal amplification (LAMP),^3,4,5^ recombinase polymerase amplification (RPA),^6,7,8^ and the integration of CRISPR-Cas systems^9,10,11,12^ which don’t require special instrumentation but can suffer from nonspecific amplification under isothermal conditions leading to false-positive results. On the other hand, protein antigen tests which work by capturing viral proteins on immobilized antibodies, such as the lateral flow assay (LFA), are less sensitive but fairly simple to use and result in a fast turnaround time.^13,14^ Nonetheless, this has led to their wide adoption as LFA tests can be performed at home without the need for bulky temperature-control or spectrophotometer instruments. Developing new platforms that combine the sensitivity of RT-qPCR with the simplicity and fast turnaround time of LFAs is needed to address current and future pandemics.

One unexplored approach for chemical sensing pertains to mechanical testing of an analyte. For example, single molecule force spectroscopy methods, such as optical tweezers^15,16^ and atomic force spectroscopy (AFM) ^17,18,19^ can identify single virus-ligand interactions with high fidelity. Thus, mechanical testing of a virus offers an alternate strategy for detection with exquisite sensitivity. Unfortunately, using force spectroscopy for analytical sensing is prohibitive because of the serial nature of these methods -interrogating one molecule at a time - and the need for expensive and dedicated instrumentation. Autonomous, force-generating motors that can be characterized in parallel may offer an alternate approach to using mechanotransduction for analytical sensing.

Herein, we present a mechanical-based viral sensing platform termed Rolosense to detect whole intact SARS-CoV-2 particles. Rolosense is a label-free and amplification-free approach which is advantageous, as such methods reduce cost and, in our case, simplify instrumentation needed for readout avoiding fluorescence dyes and spectrometers that are commonly used for nucleic acid-based assays. Our assay leverages DNA-based motors that function as the mechanical transducer reporting on specific target binding events (Fig. 1a). We leveraged our recent work using these motors to detect and transduce DNA logic operations.^20^ The motor consists of a DNA-coated spherical particle (5 μm diameter) that hybridizes to a surface modified with complementary RNA. The particle moves with speeds of >1 μm/min upon addition of ribonuclease H (RNaseH), which selectively hydrolyzes duplexed RNA and ignores single stranded RNA. The DNA motor consumes chemical energy on the RNA chip to generate piconewton mechanical work. To detect the SARS-CoV-2 virus, the DNA motors and the RNA chip are modified with virus binding ligands (i.e., aptamers) with high affinity for the S1 subunit of spike protein that is abundantly displayed on each virion.^21^ Virus binding to both the motor and surface leads to motor stalling. In other words, the microparticle moves along the surface through a “cog-and-wheel” mechanism and only specific the SARS-CoV-2 viral target acts as a “wrench” to inhibit this activity. Unlike conventional nucleic acid and protein assays that have received EUA, we do not need fluorescence nor absorbance measurements to detect a target of interest. Instead, detection of the viral target occurs when the *mechanical force* generated by the motor (∼100 pN) is insufficient to overcome the mechanical stability of the aptamer-target complex.^22^ This binding event is transduced in a label-free fashion by measuring the displacement of the motor. Another key advantage of our assay is the ability to multiplex and detect multiple respiratory viruses in the same assay. This will be critical in-patient care and in minimizing false positive results due to similar symptoms.

**Figure 1.**
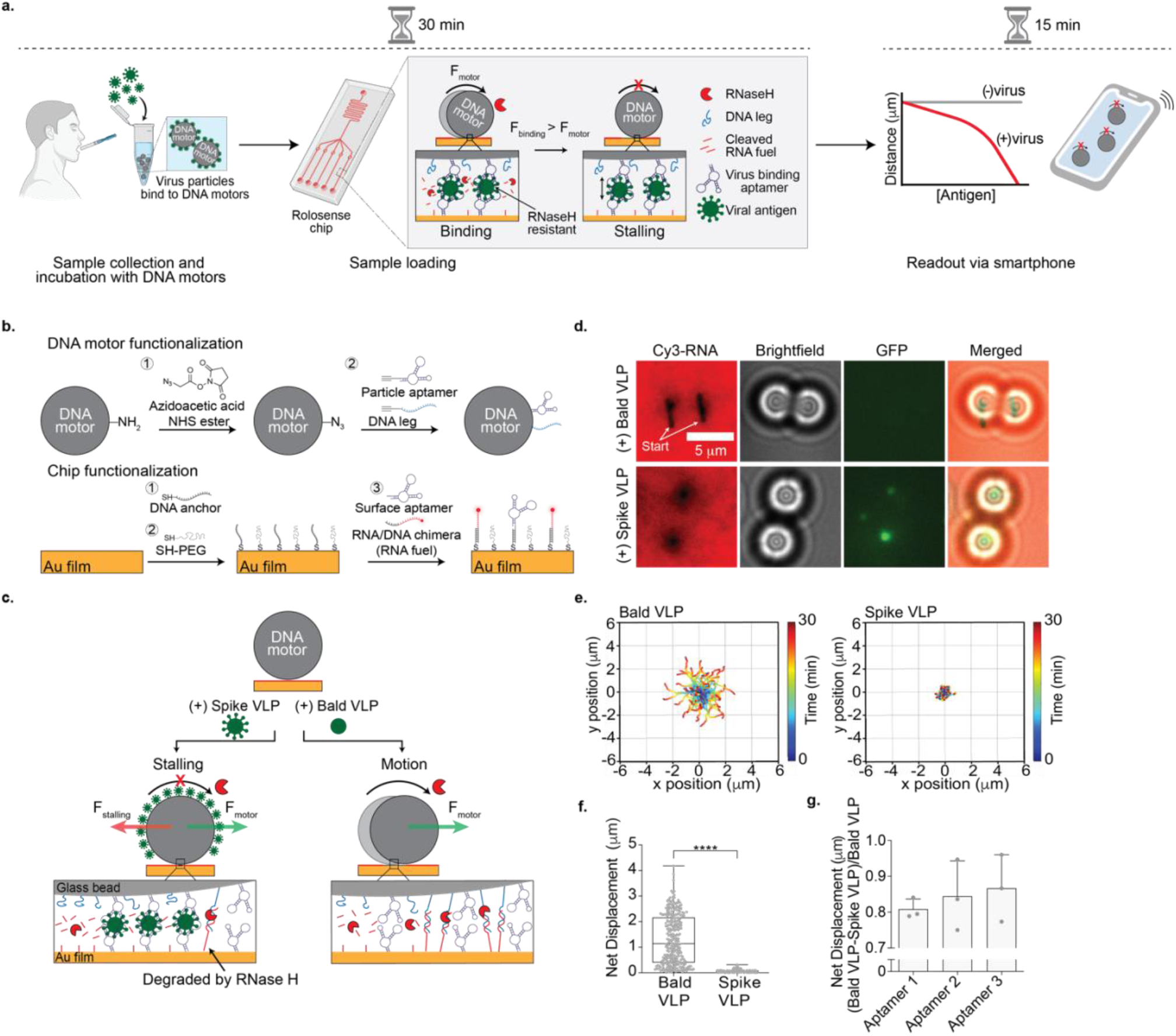
Optimizing Rolosense with GFP-labeled virus-like particles (VLPs). **a**, Schematic workflow of the Rolosense assay. The presence of virus particles leads to motor stalling which reduces the net displacement or distance travelled by the motors. Readout can be performed using simple brightfield timelapse imaging of the motors. In principle, readout can be performed in as little as 15 min using a smartphone camera. **b**, Schematic of DNA motor and chip functionalization. The DNA motors were modified with a binary mixture of with DNA leg and aptamers that have high affinity for SARS-CoV-2 spike protein. The Rolosense chip is a gold film also comprised of two nucleic acids: the RNA/DNA chimera, which is referred to as the RNA fuel, and the same aptamer as the motor. **c**, Schematic of the detection of SARS-CoV-2 virus. In the presence of VLPs expressed with spike protein (spike VLPs), the motors stall on the Rolosense chip following the addition of the RNaseH enzyme as the stalling force (red arrow) is greater than the force generated by the motor (green arrow). When incubated with the bald VLPs, or VLPs lacking the spike protein, the motors respond with motion and roll on the chip in the presence of RNAseH. **d**, Brightfield and fluorescence imaging of DNA motors detecting GFP-labeled spike VLPs. The RNA fuel was tagged with Cy3, shown here in red. Motors were incubated with 25pM of GFP-labeled bald and spike VLPs diluted in 1xPBS. Samples with GFP-labeled spike VLPs show stalled motors and no Cy3 depletion tracks in contrast to samples GFP-labeled bald VLPs. Note that stalled motors often showed GFP signal colocalization. **e**, Plots showing the trajectory of motors with bald and spike VLPs. All the trajectories are aligned to the 0,0 (center) of the plots for time = 0 min. Color indicates time (0 → 30 min). **f**, Plot showing net displacement of over 100 motors incubated with 25 pM bald and spike VLPs. **** indicates *P*<0.0001. Experiments were performed in triplicate. **g**, Plot showing the difference in net displacement between the bald/spike VLPs normalized by the bald VLP displacement in conditions using different aptamers. Each data point indicates the pooled average for an independent experiment. Error bars show the standard deviation.

Using motors modified with multivalent aptamers that have high affinity for spike protein, we were able to demonstrate a limit of detection (LoD) of 10^3^ viral copies/mL of SARS-CoV-2 WA-1, B.1.617.2, and BA.1 in artificial saliva and in exhaled breath condensate without any sample preparation or amplification steps. Note that these were the variants available to us over the course of this study. We demonstrate specific detection of SARS-CoV-2 viral particles as our motors do not respond to influenza A or other human corona viruses such as OC43 and 229E. We also show the ability to multiplex by detecting influenza A and SARS-CoV-2 in the same “pot.” Our assay can be readout *via* smartphone in as little as 15 mins. Overall, Rolosense enables rapid, sensitive, and multiplexed viral detection for disease monitoring.

## Design principles of Rolosense platform

We first functionalized DNA-based motors and chips with DNA aptamers reported in literature that had high affinity for spike protein (S1) (Supplementary Table 1 and Supplementary Fig. 1).^23,24^ Aptamers as virus binding ligands have several advantages such as ease of storage, long-term stability, and a smaller molecular weight. We started our investigations with a 50 nt S1 aptamer, aptamer 1. As depicted in Fig. 1b, the amine modified motors were functionalized and coated with a binary mixture of both the DNA leg and aptamer 1. The planar Rolosense chip was modified with a binary mixture of Cy3-labeled RNA fuel and aptamer 1. The oligonucleotides were tethered to the surface by hybridization to a monolayer of 15mer ssDNA, which we call the DNA anchor. Tuning the ratio of the DNA legs/RNA fuel to the aptamer was critical as there is a tradeoff between multivalent avidity to the virion and efficient motor motion.^20^ For example, high densities of aptamer lead to efficient virus binding but hamper processive motion. Conversely, low aptamer densities diminish virus binding, but enhance motor speed and processivity. Accordingly, we screened different ratios of aptamer/DNA leg on the particle and aptamer/RNA fuel on the chip and measured motor net displacement over a 30 min time window. We found that the introduction of aptamer at 10% density or greater on the particle or the planar surface led to a significant reduction in motor displacements (Supplementary Fig. 2). Also, motor distance was more sensitive to aptamer density on the spherical particle compared to that of the planar surface. Our results suggested that an optimal aptamer density was 10% for the particle and 50% for the planar surface, as these motors showed 1.95 +/-0.97 μm net displacement over a 30 min time window compared to the no aptamer control in which the motors traveled 2.56 +/-1.17 μm in 30 mins (Supplementary Fig. 2). Based on these results, all subsequent experiments were conducted using motors, and chips modified with 10%, and 50% aptamer density, respectively.

We first tested our assay using GFP-tagged virus-like particles (VLPs) expressing the trimeric spike protein. We used non-infectious SARS-CoV-2 S D614G HIV-1 virus-like particles (spike VLPs) (Supplementary Fig. 3). As a control to test for cross reactivity, we used GFP-tagged HIV-1 particles that lacked spike protein (bald VLPs). The motor surface was functionalized with 10% aptamer 1 and chip surface with 50% aptamer 1.^23^ The VLPs were incubated with the aptamer functionalized DNA-based motors in 1xPBS (phosphate-buffered saline) for 30 mins at room temperature. After 30 mins, the DNA-based motors were washed via centrifugation (15,000 rpm, 1 min) and then added to the Rolosense chip that was also coated with the same aptamer. In the presence of RNaseH the DNA-based motors incubated with the spike VLPs remained stalled on the surface (Fig. 1c). The VLPs were likely sandwiched between the DNA-based motor and the chip surface, and this binding led to a stalling force that halted motion. In contrast, DNA-based motors incubated with the bald VLPs lacking the spike protein translocated on the surface which is expected because the bald VLPs do not bind to the aptamers. This was confirmed by optical and fluorescence microscopy (Fig. 1d). Motors incubated with the bald VLPs displayed micron-length depletion tracks in the Cy3-RNA monolayer. The lack of fluorescence signal in the GFP channel indicates that there was minimal binding of bald VLPs to the motors. On the contrary, motors incubated with spike VLPs did not display Cy3-RNA depletion tracks and the GFP fluorescence channel showed puncta colocalized with the stalled motors confirming that the stalling was due to spike VLPs binding. Brightfield real-time particle tracking also validated this conclusion. We observed long trajectories and net displacements greater than 1.5 μm for motors incubated with bald VLPs. The spike VLP incubated motors, on the other hand, displayed short trajectories and sub 1 μm net displacements (Fig. 1e and f). Control motors without VLPs showed greater displacements than that of the bald VLP samples (Supplementary Fig. 4), likely due to non-specific bald VLPs binding. Regardless, these results demonstrate that the Rolosense design and mechanism for viral detection is valid and further motivated our subsequent experiments.

In principle, Rolosense is not unique to aptamers and virtually any virus binding ligand could be used for viral sensing. That said, in preliminary screens with two commercial antibodies, we found motor stalling with bald VLPs suggesting issues with specificity (Supplementary Fig. 5). We thus focused efforts on screening across different aptamers reported to display high affinity and specificity for SARS-CoV-2 S1. Specifically, aptamers 1, 2, and 3 ^23,24^ have reported K_D_ values in the low nanomolar range for S1. Using motors and surfaces functionalized with each of these aptamers (10% motor, and 50% chip), we showed that aptamer 3 was the most sensitive and specific for Rolosense (Fig. 1g). Based on this data, we performed all subsequent experiment using aptamer 3, unless noted otherwise.

### Detecting SARS-CoV-2 in artificial saliva

We then aimed to validate our Rolosense assay using authentic SARS-CoV-2 virus that was UV-inactivated. We tested the original SARS-CoV-2 strains first isolated in the US in the state of Washington, and hence described here as the Washington (WA-1) strain. For these sets of experiments, we spiked the virus into artificial saliva and performed Rolosense for viral readout. First, we wanted to test whether the motors and the Rolosense assay could tolerate the artificial saliva matrix since it contains mucins and divalent ions such as calcium that may interfere with the assay. Motors were suspended in artificial saliva for 30 min and then added to the aptamer-decorated chip for readout. We found that motion was not affected by the artificial saliva matrix and the motors displayed long trajectories with the addition of RNaseH enzyme and net displacement (2.20 μm +/-1.38 μm) was comparable to controls performed in 1xPBS (2.97 μm +/-1.40 μm) (Supplementary Fig. 6). Once we validated the assay in artificial saliva, we next incubated motors functionalized with 10% of aptamer 3 with 10^8^ copies/mL of SARS-CoV-2 WA-1 for 30 mins at room temperature. After 30 mins, the DNA-based motors were washed via centrifugation (15,000 rpm, 1 min) and then added to the Rolosense chip presenting aptamer 3. In the presence of RNaseH the motors remained stalled on the surface and did not display depletion tracks (Fig. 2a). Control motors without virus displayed long depletion tracks in the Cy3-RNA channel. Brightfield particle tracking confirmed these results as we observed hampered particle trajectories for SARS-CoV-2 WA-1 condition compared to the long trajectories displayed by motors without any virus (Fig. 2b). In a control experiment in which we withheld the surface aptamer, we observed that aptamer presenting motors incubated with 10^7^ copies/mL of SARS-CoV-2 WA-1 displayed long net displacements and depletion tracks in the Cy3-RNA channel (Supplementary Fig. 7). This confirmed that the stalling observed in the presence of virus is due to virus particles bridging the aptamers on the bead to the aptamers on the chip. To optimize workflow of the Rolosense assay, we tested whether we could forego the washing step following motor incubation with virus. Our results indicated that running the assay without the wash step does not degrade the integrity of the chip as the RNA on the surface remained intact (Supplementary Fig. 8). We also observed similar net displacements between the motors with and without wash when incubated with 10^7^ copies/mL of WA-1 virus. In addition, we tested whether decreasing the virus sample incubation time affects the performance of Rolosense. We show that decreasing the incubation time with the motors down to 10 minutes does not impact the performance of the Rolosense assay as the majority of the motors remained stalled (Supplementary Fig. 9).

**Figure 2.**
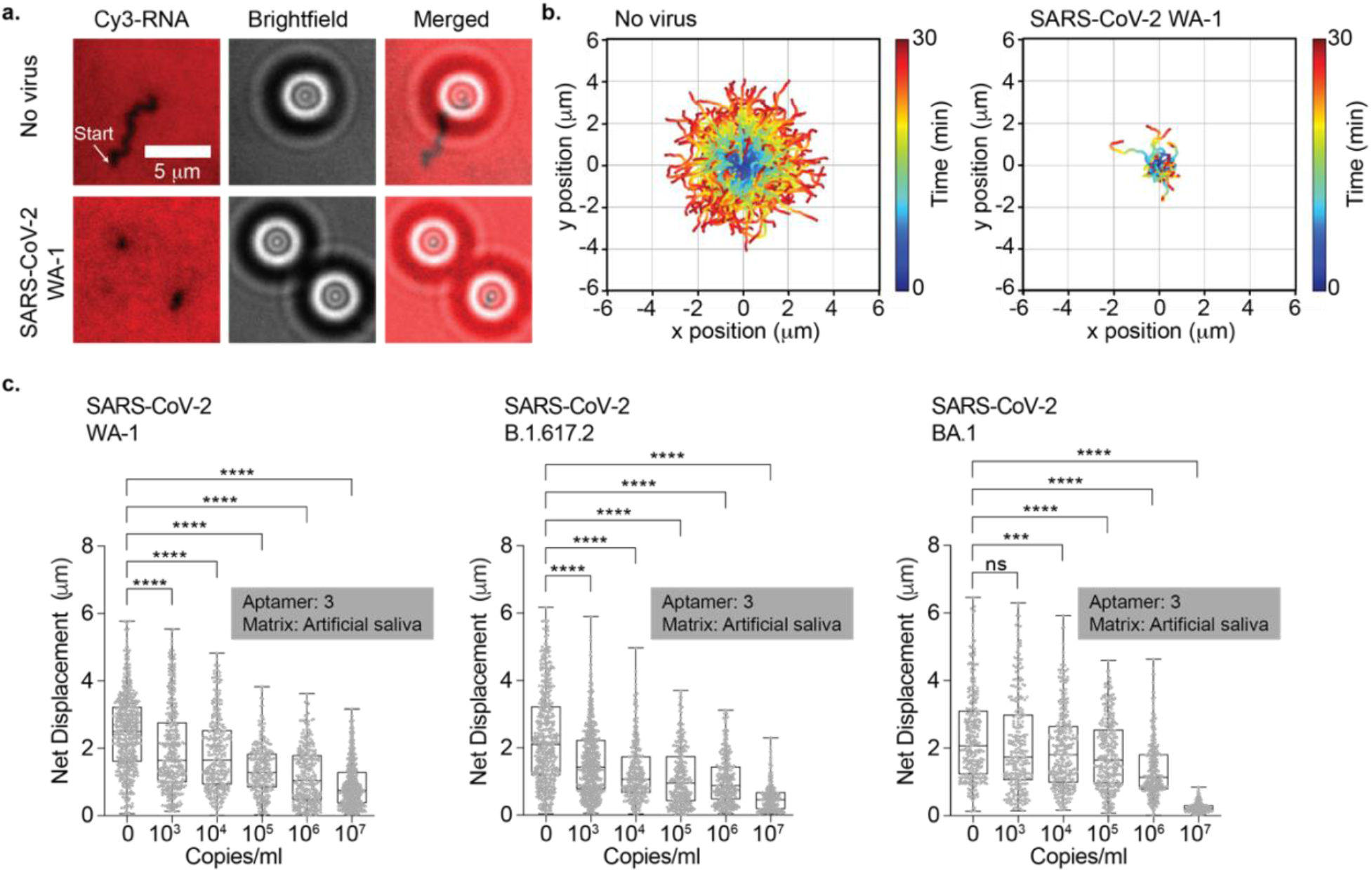
Detecting SARS-CoV-2 virus in artificial saliva. **a**, Fluorescence and brightfield imaging of DNA motors detecting the presence of 10^8^ copies/mL of UV-inactivated SARS-CoV-2 WA-1 spiked in artificial saliva. DNA motors were incubated for 30 min with the virus samples. Samples with SARS-CoV-2 show stalled motors and no depletion tracks in contrast to samples lacking the virus. **b**, Plots showing the trajectories of motors with no virus and 10^8^ copies/mL of UV-inactivated SARS-CoV-2 WA-1 strain spiked in artificial saliva. All the trajectories are aligned to the 0,0 (center) of the plots for time = 0 min. Color indicates time (0 → 30 min). **c**, Plots of net displacement of over 300 motors for each sample that was incubated with ranging concentrations of SARS-CoV-2 WA-1, B.1.617.2, and BA.1. UV-inactivated SARS-CoV-2 samples were spiked in artificial saliva and incubated with the motors functionalized with aptamer 3 at room temperature for 30 min. Each sample was performed in triplicate. **** and ns indicate *P* < 0.0001 and not statistically significant, respectively.

Next, we aimed to determine the limit of detection (LoD) of the Rolosense assay in artificial saliva. In triplicate experiments we demonstrated a LoD of ∼10^4^ copies/mL for the Washington strain (WA-1) (Fig. 2c, Supplementary Fig. 10). We also tested the Rolosense assay with other SARS-CoV-2 variants such as Delta (B.1.617.2) and Omicron (BA.1) spiked in artificial saliva. The Rolosense assay showed a sensitive response to both B.1.617.2 and BA.1 with an LoD of ∼10^3^-10^4^ copies/mL. The LoD for the B.1.617.2 and WA-1 was greater than that for the BA.1 variant which was expected given that aptamer 3 was selected using S1 of the initial Wuhan strain. Interestingly, the mutations in S1 for the B.1.617.2 strain primarily led to an increase in the net positive charge of the protein^25^ which likely aids in enhancing binding to a negatively charged aptamer. The LoD for the BA.1 strain is weaker, but this is expected given the increased number of mutations in this most recent variant. Importantly, Rolosense demonstrates an LoD that is akin to that of typical LFAs but using a DNA motor.^26,27^

To test for cross-reactivity and specificity of our assay, we measured the response of the motors incubated with other respiratory viruses such as the seasonal common cold viruses, HCoV OC43 and 229E, as well as the influenza A virus. These respiratory viruses present similar symptoms as the SARS-CoV-2 virus and thus it is important to distinguish between them. These samples were prepared in a similar manner to that of the SARS-CoV-2 variants and spiked in artificial saliva to run the Rolosense assay. As depicted in Fig. 3a, the motors displayed high specificity and responded with motion to HCoV OC43, HCoV 229E, and Influenza A which is in direct contrast to the stalling observed in the presence of SARS-CoV-2 viruses (Fig. 3b, Supplementary Fig. 11). This data, along with the LoD data, confirm that the Rolosense assay exhibits a sensitive and specific response to the SARS-CoV-2 virus which is ultimately the result of the sensitivity and specificity of aptamer 3 to its SARS-CoV-2 target.

**Figure 3.**
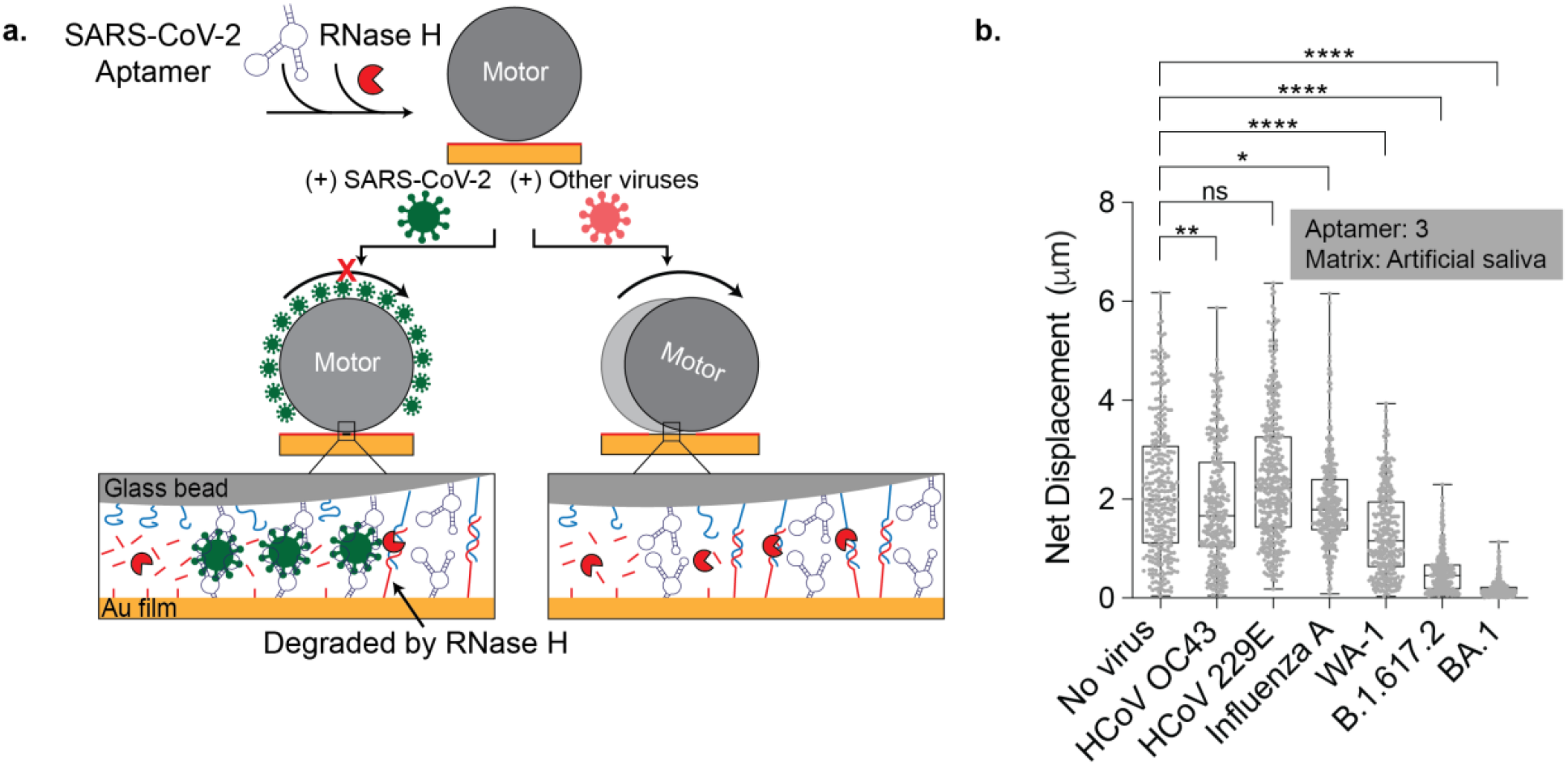
Motors demonstrate a specific response to SARS-CoV-2 viruses. **a**, Schematic of motors modified with SARS-CoV-2 aptamer stalling when incubated with SARS-CoV-2 virus particles which is in contrast to motors incubated with other viruses. **b**, Plot showing the net displacement for over 100 motors incubated with 10^7^ copies/mL of UV-inactivated HCoV OC43, HCoV 229E, influenza A, SARS-CoV-2 WA-1, SARS-CoV-2 B.1.617.2, and SARS-CoV-2 BA.1 spiked in artificial saliva. The motors were functionalized with aptamer 3 and incubated for 30 min with each sample. All measurements were performed in triplicate. ns, *, **, and **** indicate not statistically significant, *P*=0.018, *P*=0.0015, and *P*<0.0001, respectively.

### Multiplexed detection of SARS-CoV-2 and Influenza A viruses

Given the need for distinguishing between a variety of respiratory viruses, we next aimed to test whether Rolosense can detect other viruses such as the influenza A virus. This is well suited for Rolosense, as the assay is modular and can easily be programmed to detect other whole virions. We created an influenza A motor by modifying it with 10% of influenza A aptamer reported in the literature, with the chip presenting 50% aptamer.^28^ Following the protocol for SARS-CoV-2, the motors were incubated with different concentrations of influenza A virus spiked in 1xPBS for 30 min. Although the motors stalled in the presence of high concentrations of influenza A virus such as 10^10^ copies/mL, the assay performed poorly in detecting low copy numbers (Supplementary Fig. 12). To address this issue, we supplemented the 1xPBS solution with 1.5mM Mg^+2^ since divalent cations aid in secondary structure formation of aptamers.^29^ As expected, the assay improved with the addition of Mg^+2^ and we were able to detect as low as 10^4^ copies/mL of influenza A virus using this aptamer. This suggests potential for Rolosense to detect influenza A infections, as a typical swab of patients with influenza yield ∼10^8^ genome copies/ml as estimated by PCR.^30^

After validating that Rolosense is modular and can be programmed to detect other viruses, we wanted to show multiplexed detection of SARS-CoV-2 and influenza A in the “same pot.” To achieve this, we used two different motors: 5 μm silica bead functionalized with influenza A aptamer and 6 μm polystyrene bead functionalized with aptamer 3. Here we are using the size refractive index of different particles to optically encode each motor in a label free manner using brightfield contrast.^20^ The chip was functionalized with 25% influenza A aptamer and 25% aptamer 3. As depicted in Fig. 4a when the influenza A motors (5μm silica) and SARS-CoV-2 motors (6μm polystyrene) were not incubated with virus, they responded with motion in the presence of RNaseH. We observed long depletion tracks in the Cy3 channel for both motors and analysis from brightfield particle tracking of over 300 motors resulted in net displacements of 2.88 μm +/-2.00 μm and 2.68 μm +/-1.83 μm for the influenza A and SARS-CoV-2 motors, respectively (Fig. 4b, c, Supplementary movie 1). In the same tube, both motors were then incubated with 10^10^ copies/mL of the influenza A virus (in 1xPBS with 1.5mM Mg^+2^) for 30 mins at room temp. As a result, the influenza A motors remained stalled on the chip while the SARS-CoV-2 motors were free to move in the presence of RNaseH. We did not observe depletion tracks in the Cy3 channel for the influenza A motor, but the SARS-CoV-2 motors formed long depletion tracks. Brightfield particle tracking confirmed this result as the net displacement of the influenza A virus decreases, compared to no virus, and the SARS-CoV-2 motors exhibited an average net displacement of 2.60 +/-2.23 μm (Fig. 4c, Supplementary movie 2). The motors were also incubated with 10^7^ copies/mL of SARS-CoV-2 WA-1 in 1xPBS with 1.5 mM Mg^+2^. In this condition, no tracks were observed for the SARS-CoV-2 motor but the influenza A motors formed long tracks (Fig. 4b). The average net displacement of the influenza A motors was 1.97 μm +/-1.84 μm compared to 0.81 +/-0.77 μm for the SARS-CoV-2 motor (Fig. 4c, Supplementary movie 3). As a control, we also incubated the motors with both viruses, and they remained stalled on the chip (Supplementary Movie 4). All in all, using different size beads with different optical intensities we have demonstrated the possibility of multiplexed viral detection on the same chip.

**Figure 4.**
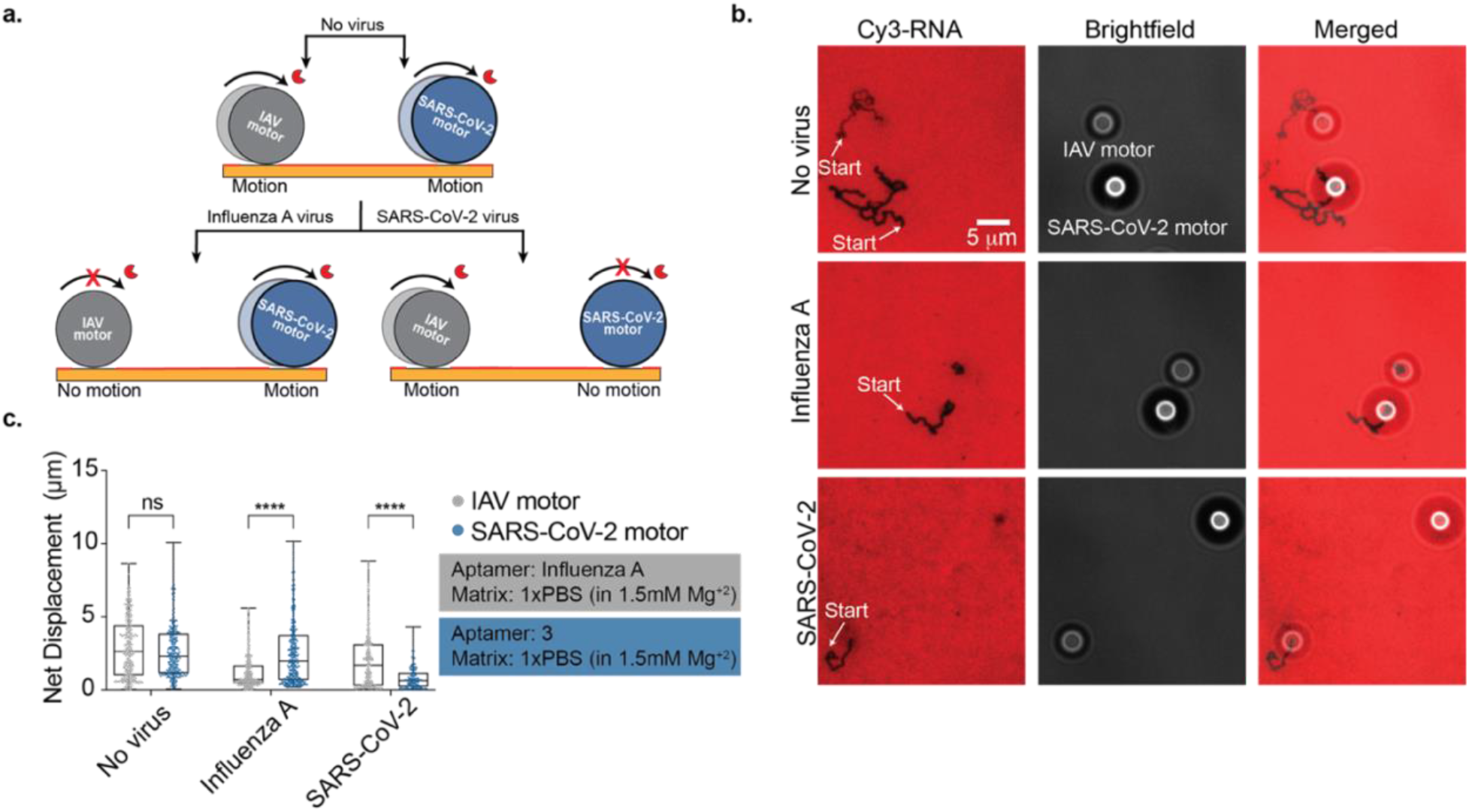
Multiplexed detection of SARS-CoV-2 and influenza A viruses. **a**, Schematic showing multiplexed detection of IAV (influenza A virus) and SARS-CoV-2. Two types of motors specific to SARS-CoV-2 (blue, 6μm polystyrene) and IAV (grey, 5 μm silica) were encoded based on the size and composition of the microparticles and used to simultaneously detect these two respiratory viruses. The two types of motors were mixed together and incubated for 30 min with the virus sample. **b**, Fluorescence and brightfield imaging of DNA motors with no virus, 10^7^ copies/mL of UV-inactivated SARS-CoV-2 WA-1, and 10^10^ copies/mL of IAV. Representative images showing the two different DNA motors are shown and each type of motor can be identified based on the brightfield particle size and contrast. Samples with SARS-CoV-2 show stalled 6 μm motors, while the IAV samples showed only stalled 5 μm particles. Samples lacking any virus showed motion of both types of motors. (**c**) Plots showing the net displacement for over 300 motors incubated with 10^7^ copies/mL of UV-inactivated SARS-CoV-2 WA-1 and 10^10^ copies/mL of IAV spiked in 1xPBS supplemented with 1.5mM Mg^+2^. Experiments were run in triplicate. ns and **** indicate not statistically significant and *P*<0.0001, respectively.

### Detecting SARS-CoV-2 *via* smartphone readout

Smartphone based sensors have captured the interest of the public health community because of their global ubiquity and their ability to provide real-time geographical information of infections.^31^ Rolosense is highly amenable to smartphone readout because smartphone cameras modified with an external lens can easily detect the motion of micron-sized motors. As a proof-of-concept we used a smartphone (iPhone 13) to detect the motion of Rolosense motors exposed to artificial saliva spiked with SARS-CoV-2. We used a simple smartphone microscope set up (Cellscope) as shown in Fig. 5a which includes an LED light source and 10x magnification lens. For these experiments, we functionalized DNA motors and chip with aptamer 3. SARS-CoV-2 WA-1 and B.1.617.2 stocks were serially diluted in artificial saliva. The DNA motors were added to these known concentrations of virus and the samples were incubated 30 mins at room temperature. Following incubation in artificial saliva, the samples were added to the Rolosense chip and imaged for motion *via* smartphone. The smartphone analyzed timelapse imaging data matched that of high-end microscopy analysis, and we found that in 15 mins timelapse videos we could detect the presence of SARS-CoV-2 in artificial saliva with an LoD of ∼10^3^ copies/mL (Fig. 5b, Supplementary movies 5 and 6). Again, matching our conclusions from high end microscopy, we observed more sensitive detection of SARS-CoV-2 B.1.617.2 than WA-1 using aptamer 3. Our results show the feasibility of label free SARS-CoV-2 sensing using smartphone camera.

**Figure 5.**
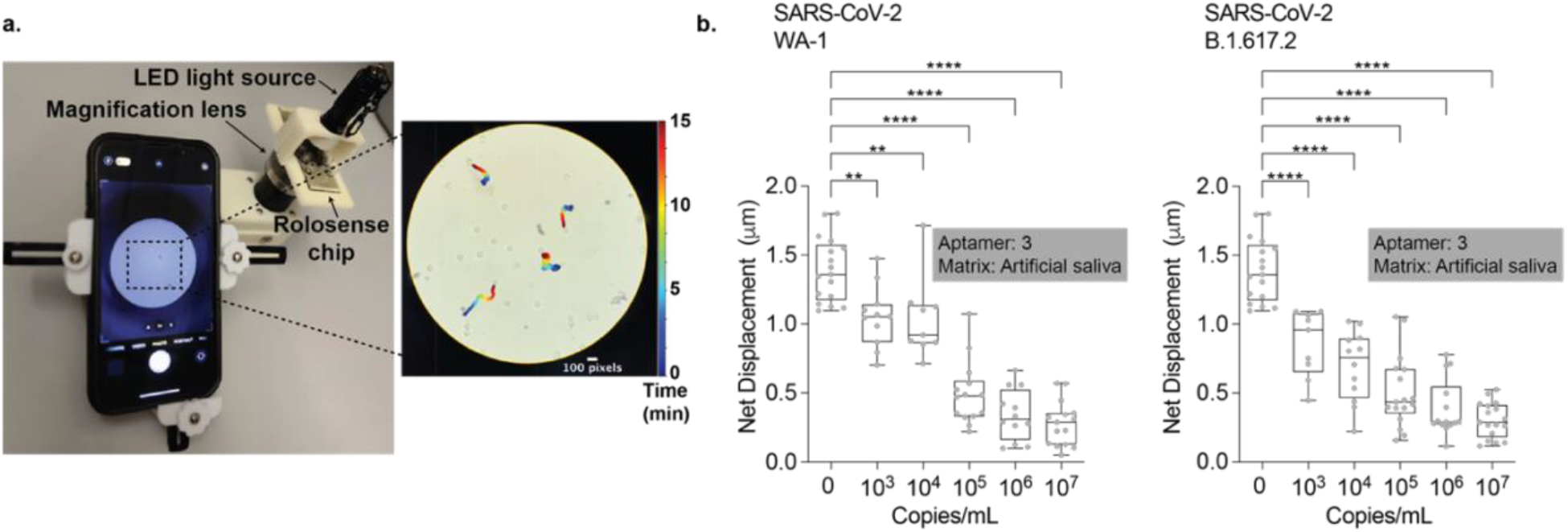
Detecting SARS-CoV-2 WA-1 and B.1.617.2 using smartphone readout. **a**, Set-up of cellphone microscope (Cellscope) which is 3D printed and includes an LED flashlight along with a smartphone holder and simple optics. The representative microscopy image shows an image of DNA motors that have been analyzed using our custom particle tracking analysis software. Moving particles show a color trail that indicates position-time (0→30 min). Scale bar is 100 pixels, and the diameter of the motors is 5 μm. **b**, Plots showing net displacement of motors incubated with different concentrations of UV-inactivated SARS-CoV-2 Washington WA-1 and B.1.617.2 samples spiked in artificial saliva. The net displacement of the motors was calculated from 15-min videos acquired using a cellphone camera. The motors were functionalized with aptamer 3 and experiments were run in triplicate. ** and **** indicate *P*=0.0015 and *P*<0.0001, respectively.

### Detecting SARS-CoV-2 in breath condensate generated samples

We aimed to better predict assay performance under real-world conditions by using exhaled breath condensate as the sensing medium since exhaled breath offers the most non-invasive and accessible biological markers for diagnosis. Exhaled breath is cooled and condensed into a liquid phase and consists of water soluble volatiles as well as non-volatile compounds.^32^ Breath condensate has already been used as a sampling media for breath analysis for detection of analytes such as viruses, bacteria, proteins, and fatty acids.^33^ We first collected breath condensate and mixed it with our motors without any virus to test whether Rolosense can tolerate breath condensate as the medium (Fig. 6). Our results indicate that breath condensate did not affect the robustness of our assay as motors without virus displayed comparable net displacements to motors diluted in 1xPBS (Supplementary Fig. 13a, b). Moreover, we also found that breath condensate displayed little DNase and RNase activity, as indicated by the high fluorescence signal of the RNA monolayer, which is well suited for Rolosense.

**Figure 6.**
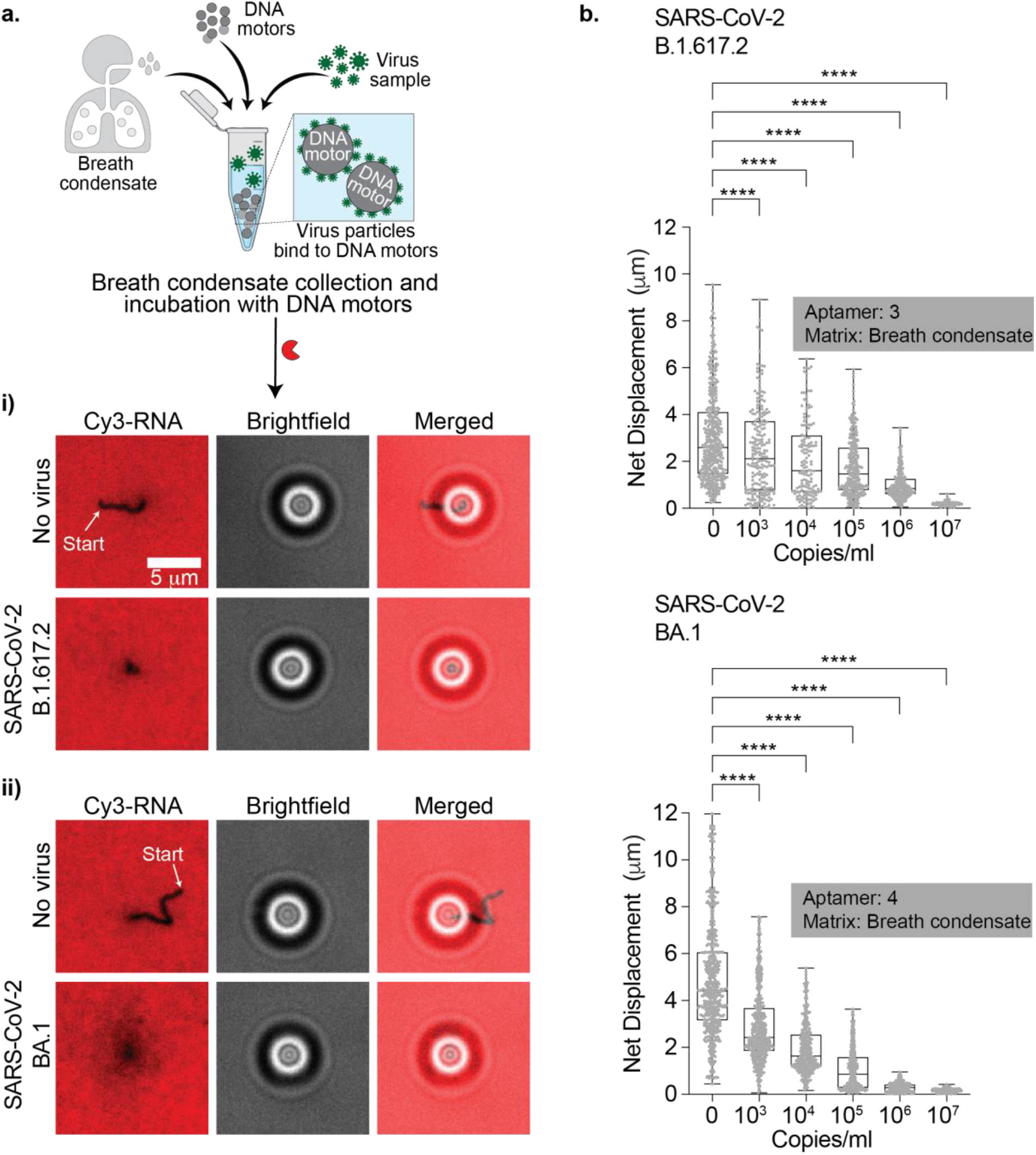
Detecting SARS-CoV-2 virus in breath condensate. **a**, (Top) Schematic of breath condensate sample collection and incubation of DNA motors with spiked-in virus particles. **i)** Fluorescence and brightfield imaging of aptamer 3 modified DNA motors without virus and with 10^7^ copies/mL of SARS-CoV-2 B.1.617.2. **ii)** Fluorescence and brightfield imaging of aptamer 4 modified DNA motors without virus and with 10^7^ copies/mL of SARS-CoV-2 BA.1. Samples without virus show long depletion tracks in the Cy3-RNA channel but no tracks are observed following sample incubation with 10^7^ copies/mL of SARS-CoV-2 B.1.617.2 and BA.1. **b**, Plots of net displacement of over 300 motors with no virus and different concentrations of UV-inactivated SARS-CoV-2 B.1.617.2, and BA.1. UV-inactivated SARS-CoV-2 samples were spiked in breath condensate and incubated with the motors functionalized with aptamer 3 (B.1.617.2) and aptamer 4 (BA.1) at room temperature. Experiments were done in triplicate. **** indicates *P* < 0.0001.

To determine the LoD of SARS-CoV-2 sensing in breath condensate, we prepared samples in a similar manner as in artificial saliva. We first functionalized the DNA-based motors and chip with aptamer 3. B.1.617.2 stocks were serially diluted in collected breath condensate. The DNA-based motors were incubated with the virus samples in breath condensate for 30 mins at room temperature. After incubation, the samples were added to the Rolosense chip and imaged. From brightfield particle tracking, we demonstrate an LoD of ∼10^3^ copies/mL for SARS-CoV-2 B.1.617.2 (Fig. 6b). Our results show that the LoD is not affected when using breath condensate as the virus sensing matrix. Over the course of this study, we became aware of a “universal” aptamer aimed at targeting the S1 subunit of the spike protein of the BA.1 variant with high affinity.^34^ Therefore to increase sensitivity in detecting the SARS-CoV-2 BA.1 variant given that it is the most widely spread variant at the moment, we functionalized our motor and chip surfaces with this BA.1 specific aptamer. As shown in Fig. 6, our data suggests that with this new aptamer we can detect as low as 10^3^ copies/mL of the BA.1 variant and possibly as low as 10^2^ copies/mL. These LoDs are highly promising as recent studies indicate that at early stages of infection with SARS-CoV-2 the estimated breath emission rate is 10^5^ virus particles/min, which suggests that 1 min of breath condensate collection will provide sufficient material for accurate SARS-CoV-2 detection.^35^

## Conclusions

We developed a mechanical-based detection method of SARS-CoV-2 viral particles that is label-free and does not require fluorescence readout or absorbance measurements. Because the motor detects the virus itself rather than the nucleic acid material, there is no need for enzymatic amplification and sample processing steps. One striking feature of Rolosense is that it represents a new concept in biosensor design in that it employs a “mechanical transduction” mechanism based on performing a mechanical test of the analyte and the outcome of this mechanical test is converting viral binding into motion output. The motors only stall if the mechanical stability of virus binding ligands, which in our case are aptamers, exceed the forces generated by the motor. The aptamer-spike protein rupture force is the fundamental parameter we are measuring rather than the Kd of the aptamer. An additional potential advantage to mechanical transduction is that it may reduce non-specific binding and detect transient interactions. While force spectroscopy has yet to be performed on aptamer-spike complexes, ACE2-spike complexes with similar affinity do show rupture forces of 57 pN when using 800 pN/s loading rates.^36^ Our past work estimated that each motor generates ∼ 100 pN of force,^22^ but likely in our case, this force is dampened because we have significantly lower density of DNA and RNA on the motor and chip, respectively. Indeed, our recent modeling^37^ suggests that lowering the magnitude of force generated by the motor can lead to enhanced biosensor performance. These estimates suggest that a single virus particle presenting 20-40 copies of trimeric spike protein^38^ will lead to motor stalling. Interestingly, when we used GFP-tagged VLPs in Rolosense we found that a population of stalled motors colocalized with single VLPs (Supplementary Fig. 14). Taken together, this suggests that Rolosense motor can respond and report on single SARS-CoV-2 virions.

We demonstrated that in artificial saliva we can detect up to 10^3^ copies/mL of SARS-CoV-2 WA-1, B.1.617.2, and the variant of concern BA.1. To validate the specificity of our Rolosense assay, we tested for cross-reactivity with other respiratory viruses such as the seasonal common cold viruses, HCoV OC43 and 229E, as well as influenza A. We did not observe a distinguishable effect on Rolosense response. A key advantage of Rolosense is the ability to multiplex and detect multiple respiratory viruses in the same assay. This capability is important for point-of-care diagnostics and in minimizing false positive results due to similar symptoms caused by other respiratory viruses. We show that by encoding different virus specific DNA motors through size and refractive index we can distinguish between SARS-CoV-2 and influenza A in one “pot.” With the aim of enabling the key steps for a point-of-care diagnostic, we demonstrated that the Rolosense motors and chip can be used to conveniently detect SARS-CoV-2 using a smartphone and a magnifying lens as the reader. The assay was performed using a rapid, ∼15 min readout without any intervention. Our assay is also suitable for exhaled breath condensate testing as we demonstrated an LoD of 10^3^ copies/mL of the B.1.617.2 and BA.1 variants.

Our reported LoD of 10^3^ copies/mL is comparable to that of lateral flow assays like the BinaxNOW COVID-19 Ag Card (Abbott Diagnostics Scarborough, Inc.) which have an LoD of 10^5^ copies/mL for the BA.1 variant.^39^ Rolosense takes advantage of multivalent binding which may contribute to LoD that is better than that of monomeric assays like LFA. Another strength of Rolosense is that it is highly modular and any whole virion that displays many copies of a target can be detected using our assay with appropriate aptamers. Also, multiplexed detection of SARS-CoV-2 and influenza A can, in principle, be scaled up to include a panel of viral targets as we could create tens of uniquely encoded motors. Such PCR panels for multiplexed detection of respiratory viruses is currently available,^40^ and hence multiplexed Rolosense would find clinical applications as our assay is rapid and can be conducted conveniently without the need for a dedicated PCR instrument.

There are two main limitations inherent to Rolosense. The first is the sensitivity to RNase and DNase contamination in the biological samples. We have documented this issue (Supplementary Fig. 15), and selective nuclease inhibitors that target RNase A, B, and C (but not RNaseH) showed excellent potential to minimize this issue. Note however, that all assays that employ DNA or RNA aptamers for biological sensing also suffer from nuclease sensitivity. Nonetheless, we were excited to see that breath condensate is relatively low in nuclease matrix and hence this is well-suited for the Rolosense assay and for detecting respiratory virions.

A second limitation is the need to employ virus binding ligands that target spike protein (or other surface displayed proteins). This is not a weakness in of itself, but rather this is a challenge when pursuing highly mutable targets such as SARS-CoV-2 and influenza that are under high evolutionary pressure to conceal their surface epitopes from immune recognition.^41^ This leads to frequent mutations in the spike protein which in contrast to nucleocapsid proteins that are infrequently altered.^42^ Our work with the “universal” spike protein aptamer shown in Figure 7 represents one solution to this problem but the specificity of universal aptamers is weaker and hence more likely to cross-react with similar spike-presenting coronaviruses. Further development and deployment of Rolosense shows potential towards a point-of-care system, which will greatly facilitate frequent, on-site molecular diagnostics.

## Methods

### Materials

All oligonucleotides were purchased from Integrated DNA Technologies (IDT), stored at 4 °C (−20 °C for RNA), and used without purification. Their sequences, including functional group modifications, are shown in Table S1. Stock solutions were made using Nanopure water (Barnstead Nanopure system, resistivity = 18.2 MΩ), herein referred to as DI water. Aminated silica beads (5 μm) were purchased from Bangs Laboratory (# SA06N). Aminated polystyrene beads (6 μm) were purchased from Spherotech (# AP-60-10). Artificial saliva was purchased from Fisher Scientific (# NC1873815). Influenza A/PR/8/34 was purchased from Charles River Laboratories (# 10100374). RNAseH was obtained from Takara Clontech (# 2150A). RTube™ breath condensate collection device was purchased from Respiratory Research (# 1025, # 3002, and # 3001). Thin Au films were generated by using a home-built thermal evaporator system. All motor translocation measurements were performed in Ibidi sticky-slide VI0.4 (Ibidi, # 80608) 17 × 3.8 × 0.4 mm channels. Smartphone microscope was obtained from Wilbur Lam, Emory University, (10×/0.25 NA objective and 20x WF eyepiece) (https://cellscope.berkeley.edu/).

### Microscopy

BF and fluorescence images were acquired on a fully automated Nikon Inverted Research Microscope Eclipse Ti2-E with the Elements software package (Nikon), an automated scanning stage, a 1.49 NA CFI Apo TIRF 100× objective, a 0.50 NA CFI60 Plan Fluor 20× objective, a Prime 95B 25mm sCMOS (scientific complementary metal-oxide semiconductor) camera for image capture at 16-bit depth, a SOLA SE II 365 Light Engine for solid state white light excitation source, and a perfect focus system used to minimize drift during timelapse. Brightfield timelapse imaging was done using 20× 0.50 NA objective with 5 sec per frame rate and an exposure time of 100 ms. Fluorescence images of Cy3 and FAM were collected using a TRITC filter set (Chroma #96321) and EGFP/FITC/Cy2/Alexa Fluor 488 Filter Set (Chroma #96226) with an exposure time of 100 ms. All imaging was conducted at room temperature.

### Viruses

UV inactivated SARS-CoV-2 and human corona (229E, OC43) virus samples at known concentrations were provided by the NIH RADx-Radical Data Coordination Center (DCC) at the University of California San Diego and BEI Resources. UV-inactivated SARS-CoV-2 Isolate hCoV-19/USA/PHC658/2021 (Lineage B.1.617.2; Delta Variant), NR-55611, was contributed by Dr. Richard Webby and Dr. Anami Patel. UV-inactivated SARS-CoV-2 Isolate hCoV-19/USA/CA-SEARCH-59467/2021 (Lineage BA.1; Omicron Variant) was contributed by Dr. Aaron Carlin and the UCSD CALM and EXCITE laboratories. Virus samples used in this study have undergone at least one freeze–thaw cycle.

### Cells and plasmids

Human embryonic kidney HEK293T/17 cell line was obtained from the ATCC (Manassas, VA, USA). Cells were grown in high glucose Dulbecco’s Modified Eagle Medium (DMEM, Mediatech, Manassas, VA, USA), 10 % Fetal Bovine Serum (FBS, Sigma, St. Louis, MO, USA), 100 U/ml penicillin–streptomycin (Gemini Bio-Products, Sacramento, CA, USA), and 0.5 mg/ml G418 sulfate (Mediatech). pCAGGS-SARS-CoV-2 S D614G (# 156421) and pcDNA3.1 vectors were obtained from Addgene (Watertown, MA, USA), and Invitrogen (Waltham, MA), respectively. HIV Gag-eGFP expression vector was a gift from Dr. Marilyn D. Resh (Memorial Sloan-Kettering Cancer Center, New York).^43^

### Virus-like particles (VLPs) production and characterization

To produce VLPs, HEK 293T/17 cells were grown to 70-80% confluency in a 100-mm plate and transfected with 4 μg SARS-CoV-2 S D614G Env, 6 μg HIV-1 Gag-eGFP, using JetPrime transfection reagent (Polyplus-transfection, Illkirch, France). For producing bald VLPs, SARS-CoV-2 S D614G expression vector was replaced with pcDNA3.1 empty vector. Twelve hours post-transfection, the media was exchanged with DMEM/10%FBS supplemented with 100 U/ml penicillin/streptomycin. At 48 h post-transfection, supernatant was collected, filtered through 0.45 μm, and precipitated with Lenti-X concentrator (Clontech) at 4 °C for 12 h. The sample was centrifuged at 4 °C, 1500 g for 45 min, the viral pellet was resuspended in 1/100 volume of PBS, aliquoted and stored at -80 °C. The number of particles was estimated based on the p24/Gag amount measured by enzyme-linked immunosorbent assay (ELISA) as previously described.^44^

### Quantification and imaging of VLPs using single particle microscopy imaging

A #1.5 glass slide (25 × 75 mm) was cleaned by sonication in DI water for 15 minutes. The sample was then sonicated in 200 proof ethanol for 15 minutes and was dried under a stream of N_2_. The glass slide was etched by piranha solution (v/v = 3:7 hydrogen peroxide/sulfuric acid, *please take caution as piranha is extremely corrosive and may expLoDe if exposed to organics*) for 30 min to remove residual organic materials and activate hydroxyl groups on the glass. The cleaned substrates were rinsed with DI water in a 200 mL beaker for 6 times and washed with ethanol three times. Slides were then transferred to a 200 mL beaker containing 2% (v/v) APTES in ethanol for 1 h, washed with ethanol three times and thermally cured in the oven (110°C) for 10 min. The slides were then mounted to 6-channel microfluidic cells (Sticky-Slide VI 0.4, Ibidi). A 1000x dilution of the GFP-tagged spike and bald VLP samples was created in 1xPBS and added to the APTES surface. High-resolution epifluorescence images (×100) of the GFP-tagged VLPs were acquired and used for further analysis.

### Thermal evaporation of gold films

A No. 1.5H ibidi glass coverslip (25 × 75 mm) (ibidi #10812) was cleaned by sonication in DI water for five minutes. The sample was then subjected to a second sonication in fresh DI water for five minutes. Finally, the slide was sonicated in 200 proof ethanol (Fischer Scientific #04-355-223) for five minutes and was subsequently dried under a stream of N_2_. The cleaned glass coverslip was then mounted into a home-built thermal evaporator chamber in which the pressure was reduced to 50 × 10^−3^ Torr. The chamber was purged with N_2_ three times, and the pressure was reduced to 1–2 × 10^−7^ Torr by using a turbo pump with a liquid N_2_ trap. Once the desired pressure was achieved, a 3 nm film of Cr was deposited onto the slide at a rate of 0.2 Å s^−1^, which was determined by a quartz-crystal microbalance. After the Cr adhesive layer had been deposited, 6 nm of Au was deposited at a rate of 0.4 Å s^−1^. The Au-coated samples were used within one week of deposition.

### Fabrication of RNA/DNA aptamer monolayers

An Ibidi sticky-Slide VI^0.4^ flow chamber was adhered to the Au-coated slide to produce six channels (17 × 3.8 × 0.4 mm dimensions). Prior to surface functionalization, each channel was rinsed with ∼5 mL of DI water. Next, thiol modified DNA anchor strands were added to each of the channels with 50 μL solution of 1 μM DNA anchor in a 1 M potassium phosphate monobasic (KHPO_4_) buffer. The gold film was sealed by Parafilm to prevent evaporation and the reaction took place overnight at room temperature. After incubation, excess DNA was removed from the channel using a ∼5 mL DI water rinse. To block any bare gold sites and to maximize the hybridization of RNA and DNA aptamer to the DNA anchoring strand, the surface was backfilled with 100 μL of a 100 μM solution of 11-Mercaptoundecyl)hexa(ethylene glycol (referred to as SH-PEG) (Sigma Aldrich #675105) solution in ethanol for six hours. Excess SH-PEG was removed by a ∼5 mL rinse with ethanol and another ∼5 mL rinse with water. For a 50% RNA and 50% DNA aptamer surface, the RNA/DNA chimera (50 nM) and the surface aptamer (50 nM) were mixed and added to the surface through hybridization to the DNA anchor in 1× PBS for 12 hours. For the multiplexed detection of SARS-CoV-2 and influenza A experiments: RNA/DNA chimera (50 nM), surface aptamer 3 (25 nM), and influenza A surface aptamer (25 nM) were mixed and added to the surface through hybridization to the DNA anchor in 1xPBS for 12 hours. The wells were again sealed with Parafilm to prevent evaporation and the resulting RNA monolayer remained stable for days.

### Synthesis of azide-functionalized motors

Before functionalization with azide, the silica and polystyrene beads were washed to remove any impurities. For the wash, 1 mg of aminated silica beads were centrifuged down for 5 minutes at 15,000 revolutions per minute (r.p.m.) in 1 mL DI water. Similarly, 1 mg of aminated polystyrene beads were centrifuged down for 10 minutes at 15,000 revolutions per minute (r.p.m.) in 1 mL DI water with 0.005% of surfactant (Triton-X). The supernatant was discarded, and the resulting particles were resuspended in 1 mL of DI water (silica beads) and 1 mL of DI water with 0.005% Triton-X (polystyrene beads). This was repeated three times and the supernatant was discarded after the final wash. Azide-functionalized particles were then synthesized by mixing 1 mg of aminated silica and polystyrene beads with 1 mg of azido acetic NHS ester (BroadPharm #BP-22467). This mixture was subsequently diluted in 100 μL of dimethylsulfoxide (DMSO) and 1 μL of a 10× diluted triethylamine stock solution in DMSO. The reaction proceeded overnight for 24 hours at room temperature and the azide-modified silica particles were purified by adding 1 mL of DI water and centrifuging down the particles at 15,000 revolutions per minute (r.p.m.) for five minutes. The azide modified polystyrene particles were purified in a similar manner except they were centrifuged for 10 minutes in 0.005% of Triton-X. The supernatant was discarded, and the resulting particles were resuspended in 1 mL of DI water. This process was repeated seven times, and during the final centrifugation step the particles were resuspended in 100 μL of DI water to yield an azide-modified particle stock. The azide-modified particles were stored at 4 °C in the dark and were used within one month of preparation.

### Synthesis of high-density DNA silica and polystyrene motors

High-density DNA-functionalized motors were synthesized by adding a total of 5 nanomoles (in 5 μL) of alkyne-modified DNA stock solution to 5 μL of azide-functionalized motors. For motors with 10% aptamer: 4.5 nanomoles of DNA leg and 0.5 nanomoles of particle aptamer 1, 2, 3, 4, or influenza A particle aptamer were mixed with 5 μL of azide-functionalized particles. The particles and DNA were diluted with 25 μL of DMSO and 5 μL of 2 M triethyl ammonium acetate buffer (TEAA). Next, 4 μL from a super saturated stock solution of ascorbic acid was added to the reaction as a reducing agent. Cycloaddition between the alkyne-modified DNA and azide-functionalized particles was initiated by adding 2 μL from a 10 mM Cu-TBTA (tris((1-benzyl-1*H*-1,2,3-triazol-4-yl)methyl)amine) stock solution in 55 vol% DMSO (Lumiprobe #21050). The reaction was incubated for 24 hours at room temperature on a shaker and the resulting DNA-functionalized motors were purified by centrifugation. The motors were centrifuged at 15,000 r.p.m. for ten minutes, after which the supernatant was discarded, and the motors were resuspended in 1 mL of a 1× PBS and 10% Triton-X (w/v) solution. This process was repeated seven times, with the motors resuspended in 1 mL 1xPBS only for the fourth to sixth centrifugations. During the final centrifugation, the motors were resuspended in 50 μL of 1xPBS. The high-density DNA-functionalized motors were stored at 4 °C and protected from light.

### Preparation of antibody coated motors and chips

To prepare DNA-antibody conjugates 25 μg of monoclonal rabbit S1 (Genetex GTX635654) and monoclonal mouse S2 (GTX632604) in 70μL 1xPBS was mixed with 80mM SMCC (succinimidyl 4-(Nmaleimidomethyl)cyclohexane-1-carboxylate) in 4 μL DMF (dimethyl formamide). The solution was incubated on ice for 2h. Excess SMCC was removed from maleimide-antibodies using Zeba spin columns (7000 MWCO, eluent: 1xPBS). Thiol-modified DNA oligo (50 nmole) were reduced using dithiothreitol (DTT, 200 mM) for 2 h at room temperature. The reduced DNA oligos were purified using NAP-5 columns (GE Healthcare). Deionized water was used as eluent. Then the reduced DNA was mixed with the maleimide-antibodies in 1xPBS overnight at 4°C. DNA-antibody conjugates were purified and concentrated using Amicon Ultra Centrifugal Filters (100 kDa MWCO). The DNA-antibody conjugates were then added to the DNA-based motors and chips via hybridization.

### Breath condensate collection

Breath condensate was collected using the R tube breath condensate collection device from Respiratory Research (# 1025, # 3002, and # 3001). The R tube breath condensate collection device consists of three parts: the disposable R tube collector, the cooling sleeve, and the plunger. First, the cooling sleeve was placed in a -20°C freezer for 15 mins. After 15 mins, the cooling sleeve was placed on top of the disposable R tube collector and exhaled breath condensate was collected by breathing into the mouthpiece of the R tube collector for 2-5 mins. The vapor emerging from the breath was condensed onto the sides of the R tube collector. Following 2-5 mins of breathing into the R tube collector, the mouthpiece was removed from the bottom of the R tube collector and the tube was placed on top of the plunger and pushed through it. The exhaled breath condensate was collected into a pool of liquid at the top. The condensed breath was then transferred into an Eppendorf tube and used in creating serial dilutions of the virus samples.

### Motor translocation

Before beginning the experiments, known concentration of virus samples were serially diluted in either 1xPBS, artificial saliva, or collected breath condensate to create samples of different virus concentrations. The DNA-based motors were then incubated with different concentrations of virus samples for 30 mins at room temperature. This was done by adding 1 μL of DNA-based motors (∼800 particles/μL) in 49 μL of either 1xPBS (+/-virus particles) or matrix such as artificial saliva or breath condensate (+/-virus particles). After 30 mins of incubation, the DNA-based motors were added to the Rolosense chip which was pre-washed with 5 mL of 1 x PBS to remove excess unbound RNA and surface aptamer. Motor translocation was then initiated with 100 μL rolling buffer which consisted of 73 μL DI water (73%), 5 μL of 10x RNaseH reaction buffer (25 mM Tris pH 8.0, 8 mM NaCl, 37.5 mM KCl, 1.5 mM MgCl_2_), 10 μL of formamide (10%), 10 μL of 7.5% (g ml^−1^) Triton X (0.75%), 1 μL of RNaseH in 1 × PBS (five units), and 1 μL of 1 mM DTT (10 μM). RNAseH enzyme stock was stored on ice for up to 2 hours. Particle tracking was achieved through BF imaging by recording a timelapse at five second intervals for 30 minutes via the Nikon Elements software. High-resolution epifluorescence images (×100) of fluorescence-depletion tracks as well as VLP fluorescence intensity were acquired to verify that particle motion resulted from processive RNA hydrolysis and confirm VLP binding. The resulting timelapse files and high-resolution epifluorescence images were then saved for further analysis.

### Image processing and particle tracking

Image processing and particle tracking was performed in Fiji (ImageJ) as well as python. Timelapse app *Lapse It* v. 5.02 was used to record timelapse videos (5 sec/frame) of DNA motors on smartphone. The bioformats toolbox in Fiji (ImageJ) enabled direct transfer of Nikon Elements image files (*.nd2) into the Fiji (ImageJ) environment where all image/video processing was performed. Particle tracking was performed using the 2D/3D particle tracker from the Mosaic plugin in Fiji (ImageJ)^45^ in which we generated .csv files with particle trajectories that were used for further analysis. The algorithms for processing the data for motor trajectories, net displacements, and speeds were performed on python v. 3.7.4. Calculation of drift correction was adapted from trackpy (*github*.*com/softmatter/trackpy*). Full python script from brightfield acquisition data can be found at https://github.com/spiranej/particle_tracking_. Statistical analyses were performed in GraphPad v. 9.1.0

## Data availability

Source statistical data are provided with this paper. Additional data sets generated are available from the corresponding author on reasonable request.

## Code availability

Python script from brightfield acquisition data regarding net displacements and particle ensemble trajectories can be found at https://github.com/spiranej/particle_tracking_.

## Acknowledgements

We acknowledge support from NIH grant U01AA029345-01, NSF DMR 1905947, NSF MSN 2004126, and NSF PHY 1806833. We thank Suzie Pun for helpful discussions on the aptamers used for SARS-CoV-2 sensing. We thank Swaminathan Rajaraman and Frank Sommerhage for helpful discussions regarding the design of Rolosense chip and reader. We also thank Sergey Urazhdin for access to the thermal evaporator and Wilbur Lam for Cellscope.

## Author Contributions

S.P. and K.S. conceived the project. S.P. designed all experiments, analyzed data, and compiled the figures. L.Z. helped with the preparation of the Rolosense chips. A.B. helped with the microscopy imaging of the VLPs and related discussions. M.M. and G.B.M. produced the VLPs that were used in the proof-of-concept experiments and helped with related discussions. S.P. and K.S. wrote the manuscript. All authors helped revise the manuscript.

