## Supplementary Information for "Rolosense: Mechanical detection of SARS-CoV-2 using a DNA-based motor"

**Supplementary Table 1: Oligonucleotide sequence design**

| ID | Sequences (5'-3') |
|---|---|
| DNA Anchor | /5AmMC6/GAGAGAGATGGGTGCTTTTTTTTTTTTTTT/3ThiolMC3-D/ |
| RNA/DNA Chimera | GCACCCATCTCTCTCCCCCCrCrUrGrUrGrArUrUrGrArUrUrArCrU/3Cy3Sp/ |
| DNA Leg | /5Hexynyl/TTTTTTTTTTTTTTTAGTAATCAATCACAG |
| Particle Aptamer 1[1] | /5Hexynyl/TTTTTCGCGGTCATTGTGCATCCTGACTGACCCTAAGGTG CGAACATCGCCCGCG |
| Surface Aptamer 1 | CGCGGTCATTGTGCATCCTGACTGACCCTAAGGTGCGAACATCGCC CGCGGCACCCATCTCTCTC |
| Particle Aptamer 2[2] | /5Hexynyl/TTTTTGGGAGAGGAGGGAGATAGATATCAACCCATGGTAG GTATTGCTTGGTAGGGATAGTGGGCTTGATGTTTCGTGGATGCCACA GGAC |
| Surface Aptamer 2 | GGAGAGGAGGGAGATAGATATCAACCCATGGTAGGTATTGCTTGGTA GGGATAGTGGGCTTGATGTTTCGTGGATGCCACAGGACGCA CCC AT CTCTCTC |
| Particle Aptamer 3[2] | /5Hexynyl/TTTTTCCCATGGTAGGTATTGCTTGGTAGGGATAGTGGG |
| Surface Aptamer 3 | CCCATGGTAGGTATTGCTTGGTAGGGATAGTGGGGCACCCATCTCTCTC |
| Particle Aptamer 4[3] | /5Hexynyl/TTTTTACGCCAAGGTGTCACTCCGTAGGGTTTGGCTCCGGGCC TGGCGTCGGTCG CGAAGCATCT CCTTGGCGT |
| Surface Aptamer 4 | ACGCCAAGGTGTCACTCCGTAGGGTTTGGCTCCGGGCCTG GCGTCGGTCGCGAAGCATCTCCTTGGCGTGCACCCATCTCTCTC |
| Influenza A Particle[4] Aptamer | /5Hexynyl/TTTTTGGCAGGAAGACAAACAGCCAGCGTGACAGCGACGCG TAGGGACCGGCATCCGCGGGTGGTCTGTGGTGCTGT |
| Influenza A Surface Aptamer | GGCAGGAAGACAAACAGCCAGCGTGACAGCGACGCGTAGGGACCGGC ATCCGCGGGTGGTCTGTGGTGCTGTGCA CCC ATC TCT CTC |

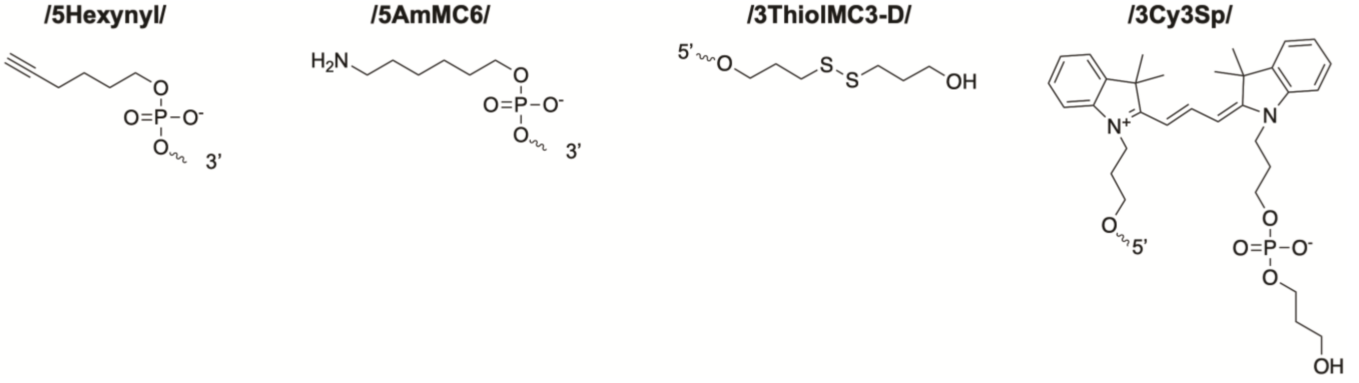


**Supplementary Table 1.** Table summarizing the sequences of oligonucleotides used in the design of the Rolosense assay displayed in a 5' to 3' orientation. The 5' and 3' DNA and RNA modifications are indicated in the table and illustrated below it.

**Figure S1. Secondary structure prediction and alignment of aptamers.**

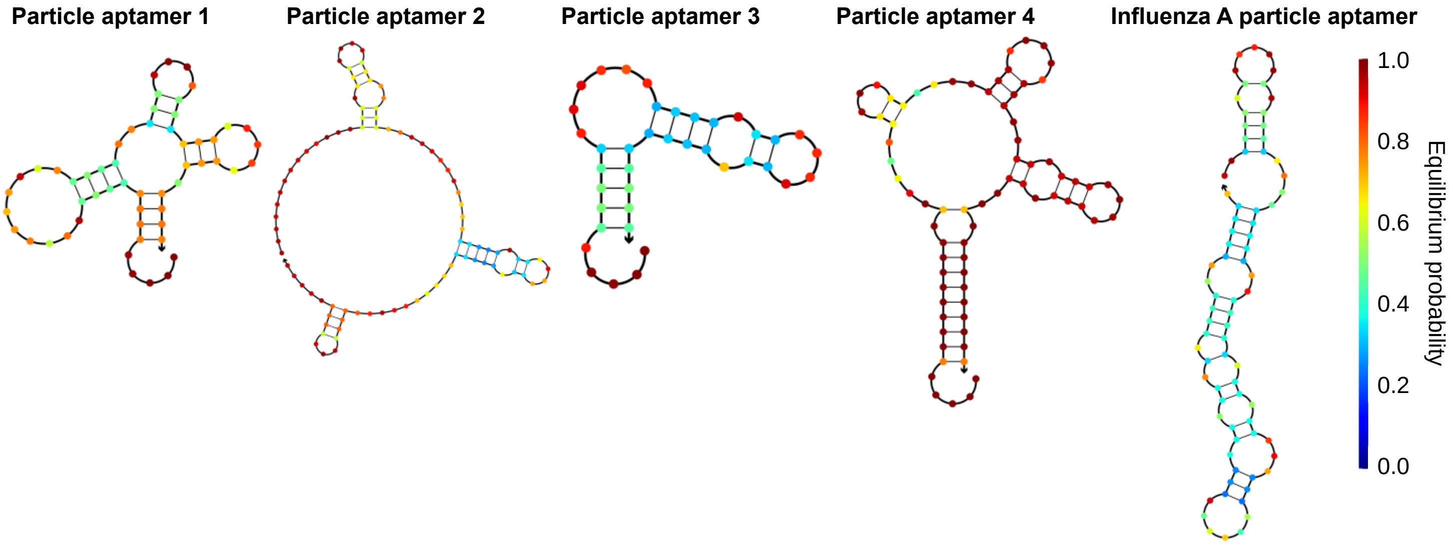


**Figure S1. Secondary structure prediction and alignment of aptamers.** (**a**) Secondary structures were predicted using NUPACK (http://www.nupack.org/) for aptamers 1, 2, 3, 4 and influenza A aptamer with the following conditions: 25˚C, 0.137 M $Na^+$. Arrows represent the 3' end of the aptamer. Nucleotides are color-coded by their equilibrium probability (legend on right).

**Figure S2. DNA-based motors tolerate certain densities of aptamer.**

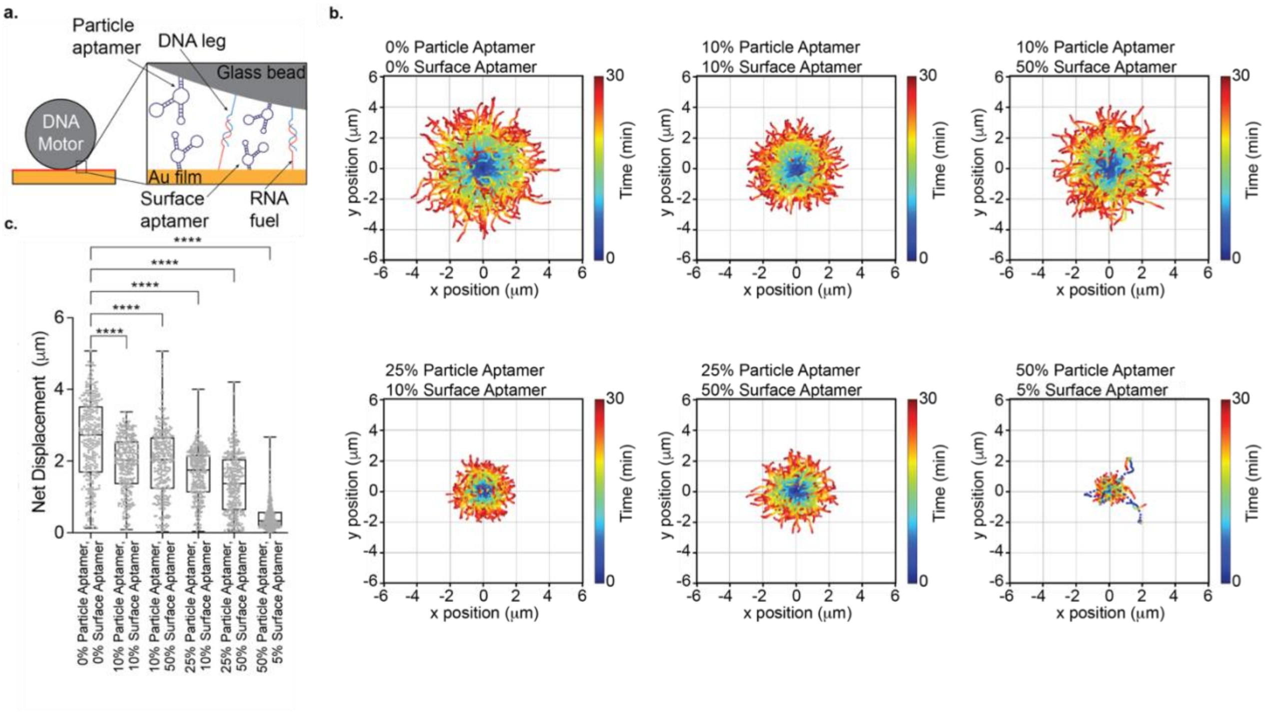


**Figure S2. DNA-based motors tolerate certain densities of aptamer. a,** Schematic of DNA motor and chip comprised of particle aptamer, DNA leg, surface aptamer, RNA fuel respectively. DNA motors are modified with a binary mixture of DNA leg and particle aptamer. The Rolosense chip is modified with a binary mixture of RNA fuel and surface aptamer. **b,** Plots showing the trajectory of motors with different densities of particle and surface aptamer 1. All the trajectories are aligned to the 0,0 (center) of the plots for time = 0 min. Color indicates time (0 → 30 min). **c,** Plots of net displacement of ~300 motors with 0% particle aptamer, 0% surface aptamer; 10% particle aptamer, 10% surface aptamer; 10% particle aptamer, 50% surface aptamer; 25% particle aptamer, 10% surface aptamer; 25% particle aptamer, 50% surface aptamer; 50% particle aptamer, 5% surface aptamer. 0% particle aptamer and 0% surface aptamer serves as the control with only DNA leg and RNA fuel. **** indicates $P < 0.0001$.

**Figure S3. Quantification of VLPs using single particle imaging.**

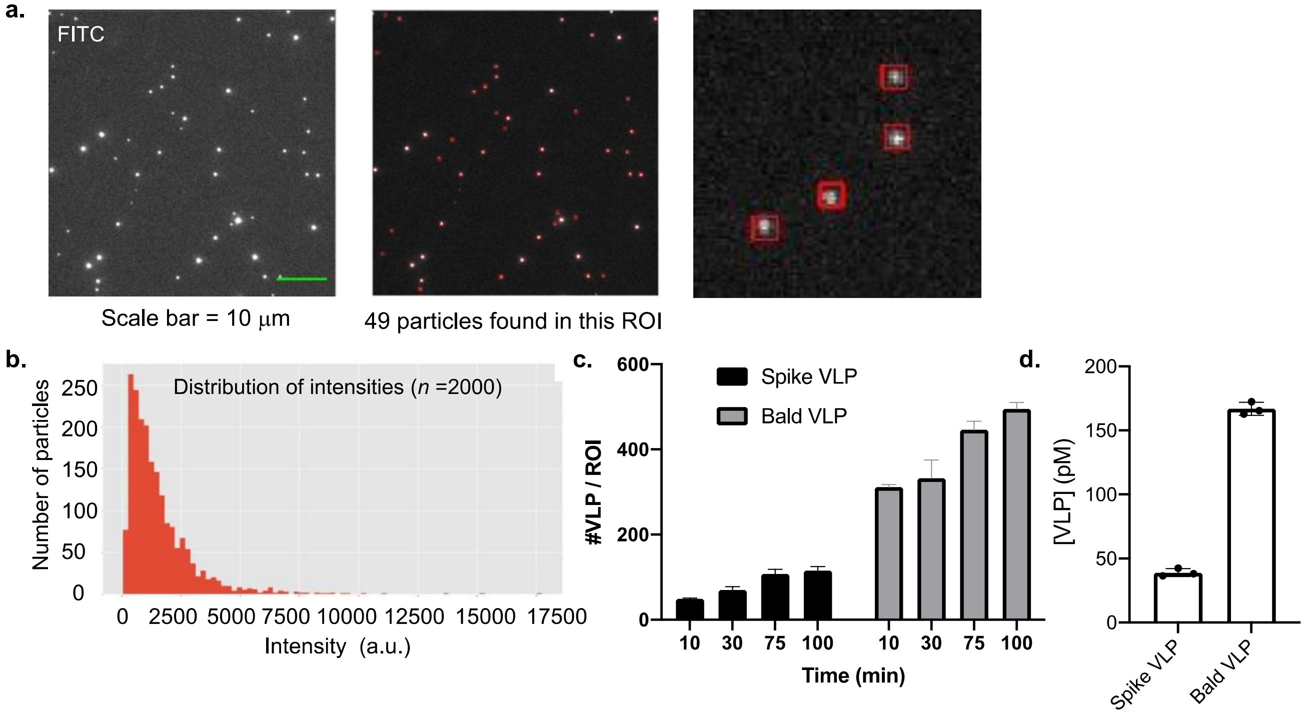


**Figure S3. Quantification of VLPs using single particle imaging. a,** Fluorescence imaging of VLPs on an amine modified glass slide. **b,** Distribution of the fluorescence intensity of 2,000 particles. **c,** Quantification of VLPs per ROI to determine density of particles per $\mu m^2$. **d,** Determined concentration of stock VLP samples which was obtained using the area of the imaging channel ($6.46 \times 10^7$ $\mu m^2$) and total volume of VLP samples (100 μL): $\frac{\#\ of\ VLPs\ per\ ROI * 6.46\text{x}10^7\ \text{um}^2}{100\ \text{uL}}$.

**Figure S4. Net displacement of aptamer-modified motors without VLPs.**

**a.**

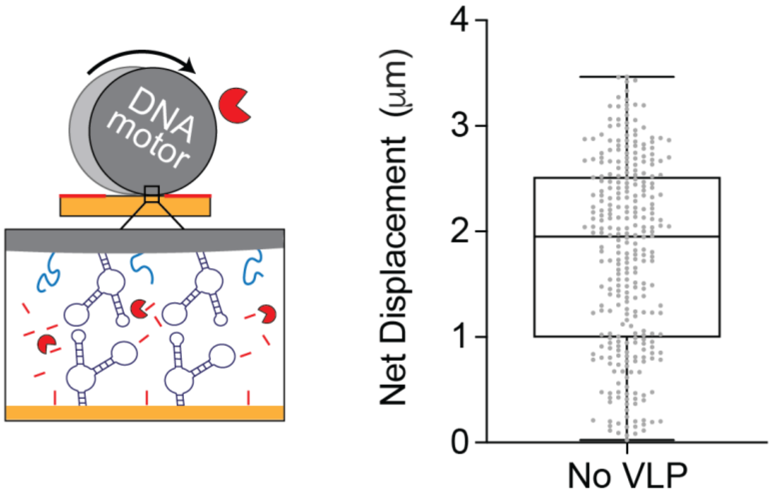


**Figure S4. Net displacement of aptamer-modified motors without VLPs. a,** (Left) Schematic of DNA motor and chip functionalized with aptamer 1. (Right) Plot showing net displacement of over 100 motors with no VLP. Experiments were done in triplicate and the net displacement for these motors was 1.87 μm +/- 0.88 μm.

**Figure S5. Using motors modified with antibodies to detect SARS-CoV-2.**

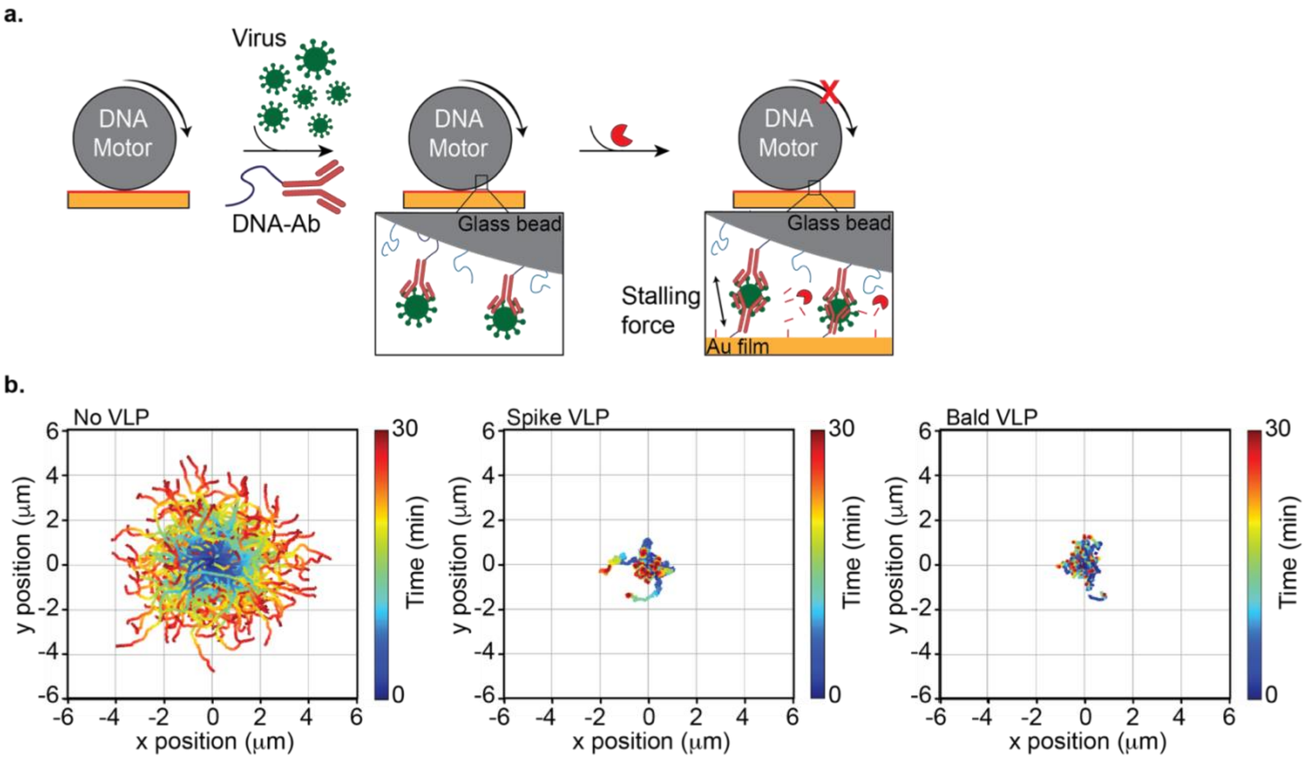


**Figure S5. Using motors modified with antibodies to detect SARS-CoV-2. a,** Schematic of Rolosense detection of SARS-CoV-2 virus particles using a commercially available antibody with high affinity for S1 subunit of spike protein. The DNA-S1 antibody conjugate was added to the motors and chip via hybridization. **b,** Plots showing the trajectory of motors without VLP, 25pM spike VLP, and 25pM bald VLP. All the trajectories are aligned to the 0,0 (center) of the plots for time = 0 min. Color indicates time (0 → 30 min). The DNA motors modified with antibodies did not show a specific response to the spike protein detection.

**Figure S6. DNA-based motors tolerate artificial saliva as a medium for virus sensing.**

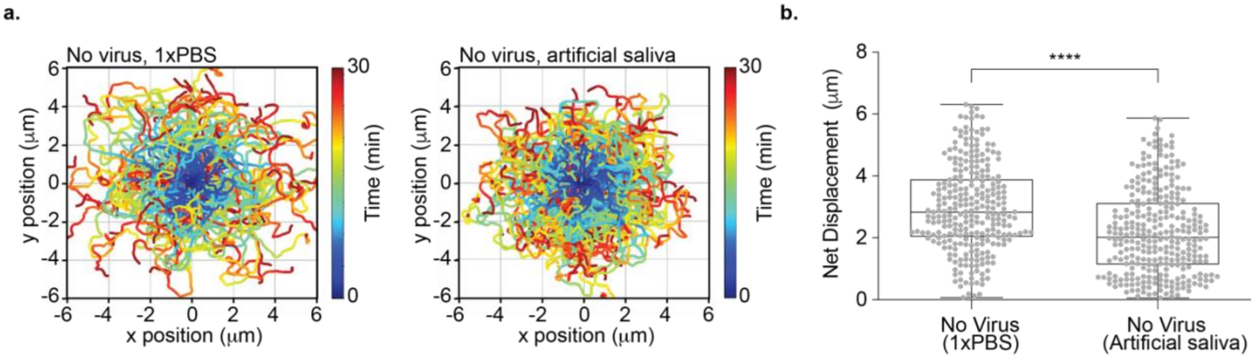


**Figure S6. DNA-based motors tolerate artificial saliva as a medium for virus sensing. a,** Plots showing the trajectory of motors without virus diluted in 1xPBS (phosphate-buffered saline) and artificial saliva. All the trajectories are aligned to the 0,0 (center) of the plot for time = 0 min. Color indicates time (0 → 30 min). The motors were modified with 10% aptamer 3 and the chip with 50% aptamer 3. **b,** Plots of net displacement of ~300 motors without virus diluted in 1xPBS and artificial saliva. **** indicates $P < 0.0001$. Experiments were performed in triplicate.

**Figure S7. DNA-based motors incubated with virus bind specifically to aptamers on the Rolosense chip.**

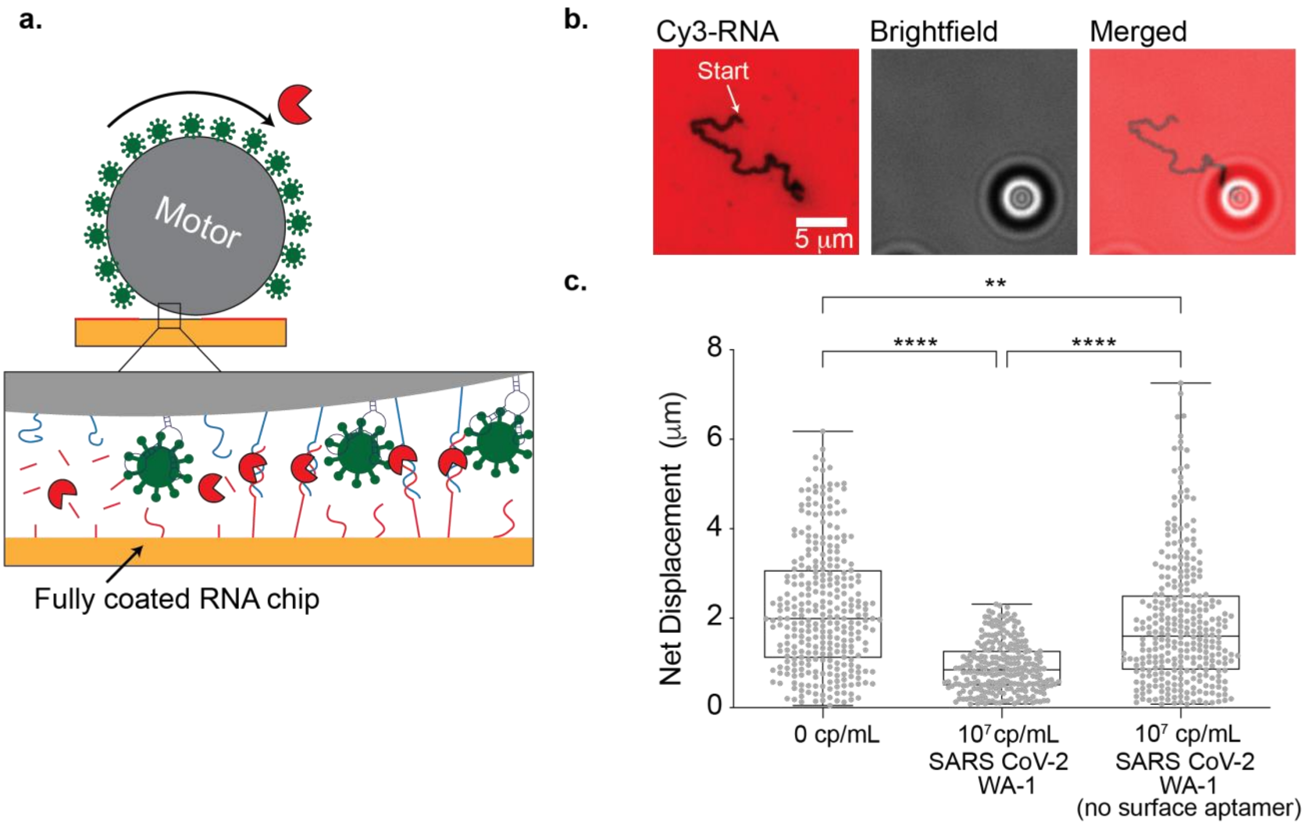


**Figure S7. DNA-based motors incubated with virus bind specifically to aptamers on the Rolosense chip.** **a,** Schematic of DNA motor functionalized with aptamer 3 rolling on a fully coated RNA chip (no aptamer) in the presence of the RNaseH. **b,** Fluorescence and brightfield imaging of DNA motors modified with 10% aptamer 3 incubated with $10^7$ copies/mL of UV-inactivated SARS-CoV-2 WA-1 spiked in 1xPBS for 30 mins at room temperature. The motors formed long RNA depletion tracks on the chip that was not functionalized with surface aptamer. The RNA fuel was tagged with Cy3, shown here in red. **c,** Plots of net displacement of ~300 motors with 10% particle aptamer, 50% surface aptamer with no virus; 10% particle aptamer, 50% surface aptamer with $10^7$ copies/mL of UV-inactivated SARS-CoV-2 WA-1; and 10% particle aptamer, 0% surface aptamer with $10^7$ copies/mL of UV-inactivated SARS-CoV-2 WA-1. ** and **** indicate $P$=0.0015 and $P$ <0.0001, respectively. Experiments were performed in triplicate.

**Figure S8. Eliminating washing steps does not affect the integrity of Rolosense.**

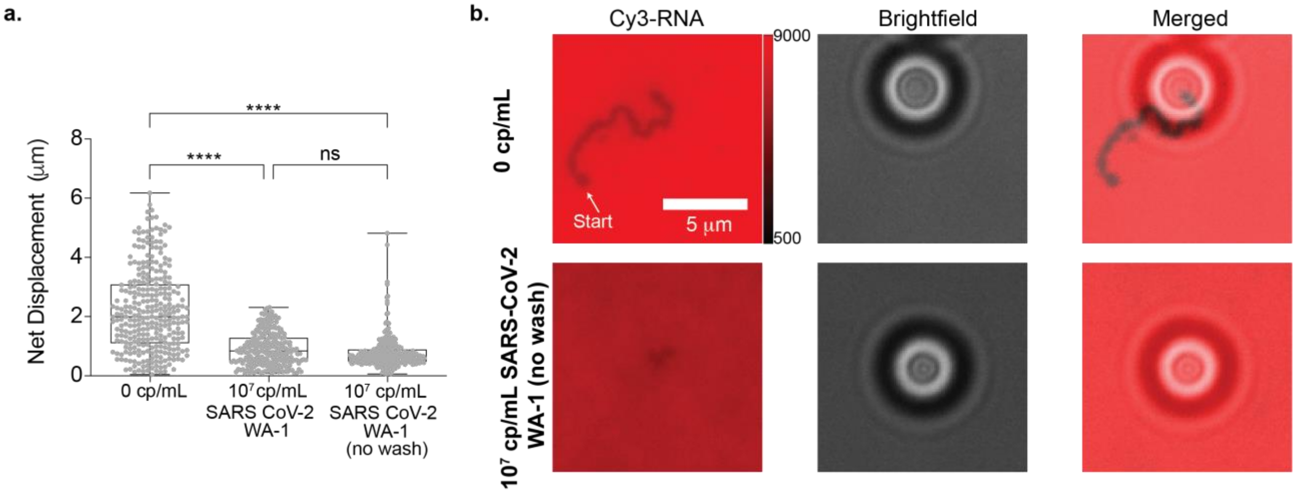


**Figure S8. Eliminating washing steps does not affect the integrity of Rolosense. a,** Plots of net displacement of ~300 motors with no virus, $10^7$ copies/mL of UV-inactivated SARS-CoV-2 WA-1 in artificial saliva, and $10^7$ copies/mL of UV-inactivated SARS-CoV-2 WA-1 in artificial saliva with no washing step following virus incubation with the DNA motors. **** and ns indicate $P < 0.0001$ and not statistically significant, respectively. Experiments were performed in triplicate. **b,** Fluorescence and brightfield imaging of DNA motors without virus diluted in artificial saliva. The motors form depletion tracks in the Cy3-RNA channel in the presence of RNaseH. When the DNA motors are incubated with $10^7$ copies/mL of SARS-CoV-2 WA-1 spiked in artificial saliva without the washing step, they do not form depletion tracks in the Cy3-RNA channel. However, foregoing the wash after sample incubation does not compromise the integrity of RNA on the chip and Rolosense continues to function. The vertical color bar indicates the minimum and maximum values (a.u.) used to display the fluorescence of the Cy3-tagged RNA. We include this bar to note the integrity of the RNA monolayer.

**Figure S9. Decreasing DNA motor and virus incubation time does not affect the overall performance of Rolosense.**

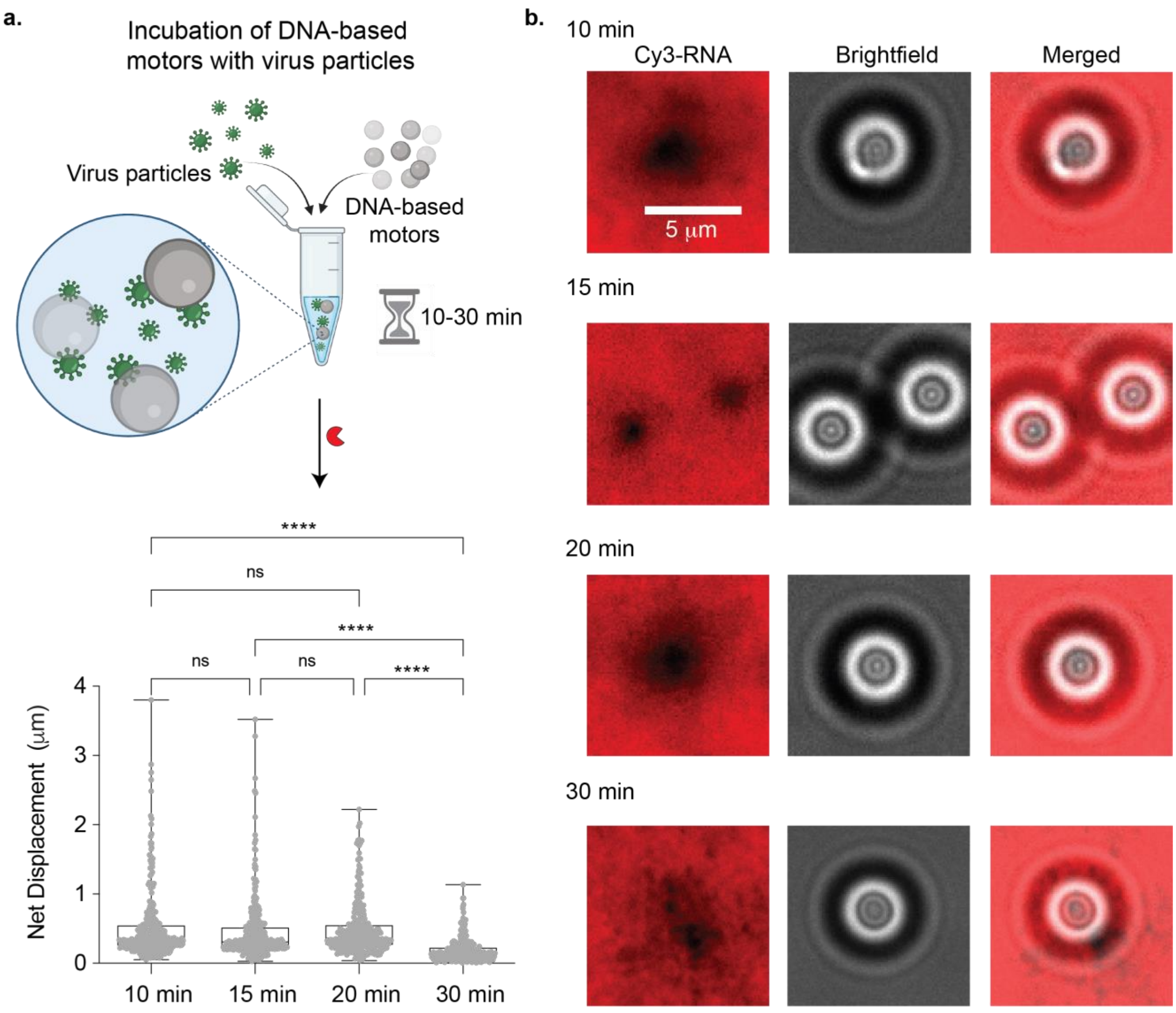


**Figure S9. Decreasing DNA motor and virus incubation time does not affect the overall performance of Rolosense.** **a,** (Top) Schematic of sample generation which includes incubation of virus with DNA motors ranging from 10 to 30 mins. (Bottom) Plots of net displacement of ~300 motors with $10^7$ copies/mL of UV-inactivated SARS-CoV-2 BA-1 in 1xPBS at different sample incubation time points following RNAseH addition. ns and **** indicate not statistically significant and *P* < 0.0001, respectively. Experiments were performed in triplicate. **b,** Fluorescence and brightfield imaging of DNA motors incubated with $10^7$ copies/mL of SARS-CoV-2 BA.1 at different sample incubation time points. No depletion tracks are observed for samples incubated at 10 min, 15 min, 20 min, or 30 min.

**Figure S10. Detection sensitivity of SARS-CoV-2 in artificial saliva.**

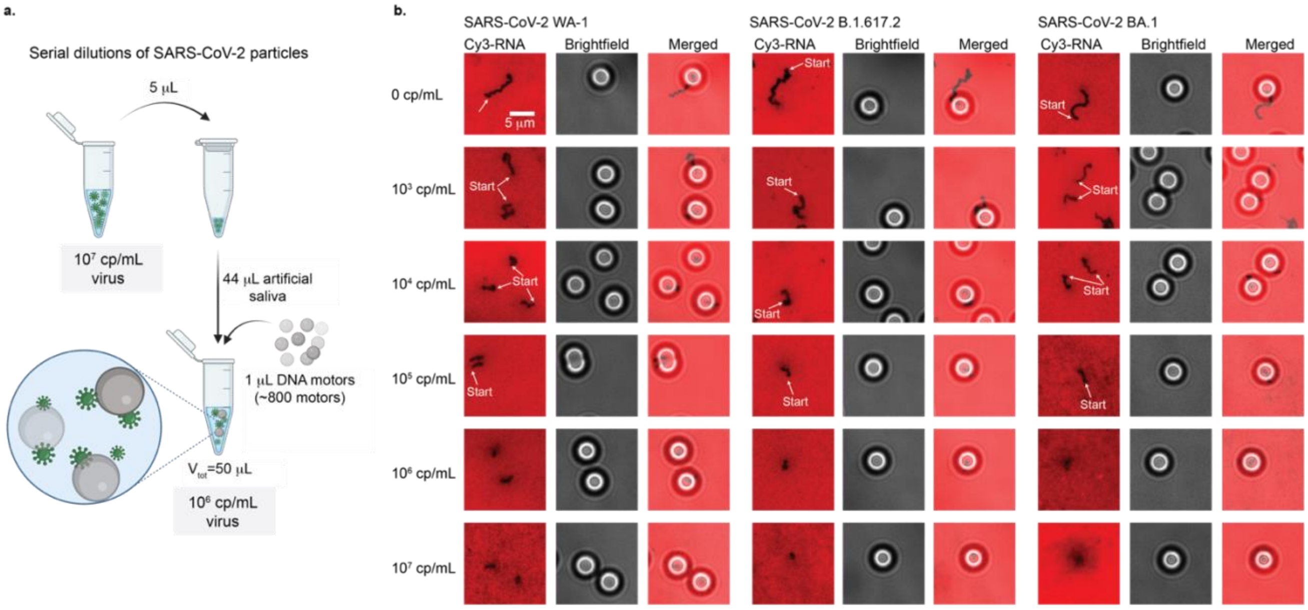


**Figure S10. Detection sensitivity of SARS-CoV-2 in artificial saliva. a,** Schematic of artificial saliva sample generation. Serial dilutions of SARS-CoV-2 particles are created in a 50 µL total volume which includes 44 µL of artificial saliva and 1 µL of DNA motors (~800 motors). **b,** Fluorescence and brightfield imaging of DNA motors incubated with different concentrations of UV-inactivated SARS-CoV-2 WA-1, B.1.617.2, and BA.1 spiked in artificial saliva. For all three variants, the DNA motors display decreasing depletion track length in the Cy3-RNA channel as virus concentration increases.

**Figure S11. Motors demonstrate a specific response to SARS-CoV-2 viruses.**

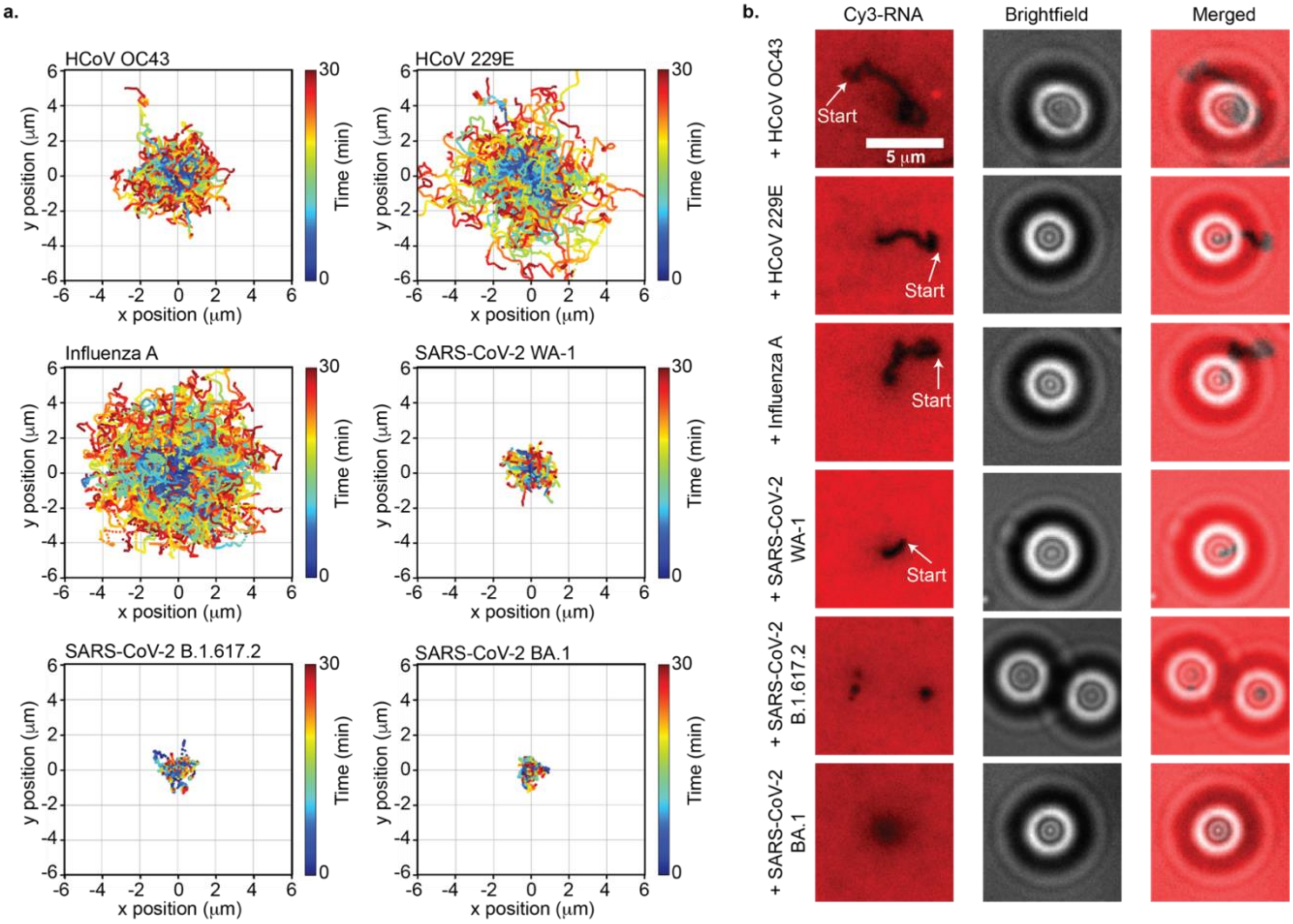


**Figure S11. Motors demonstrate a specific response to SARS-CoV-2 viruses. a,** Plots showing the trajectory of motors incubated with $10^7$ copies/mL of UV-inactivated HCoV OC43, HCoV 229E, influenza A, SARS-CoV-2 WA-1, B.1.617.2, and BA.1 in artificial saliva. All the trajectories are aligned to the 0,0 (center) of the plots for time = 0 min. Color indicates time (0 → 30 min). **b,** Fluorescence and brightfield imaging of DNA motors with $10^7$ copies/mL of UV-inactivated HCoV OC43, HCoV 229E, influenza A, SARS-CoV-2 Washington WA-1, SARS-CoV-2 B.1.617.2, and SARS-CoV-2 BA.1 spiked in artificial saliva. Motors incubated with UV-inactivated SARS-CoV-2 WA-1, B.1.617.2, and BA.1 showed no RNA depletion tracks whereas motors incubated with UV-inactivated HCoV OC43, HCoV 229E, and influenza A formed long RNA depletion tracks.

**Figure S12. DNA motors detect influenza A virus.**

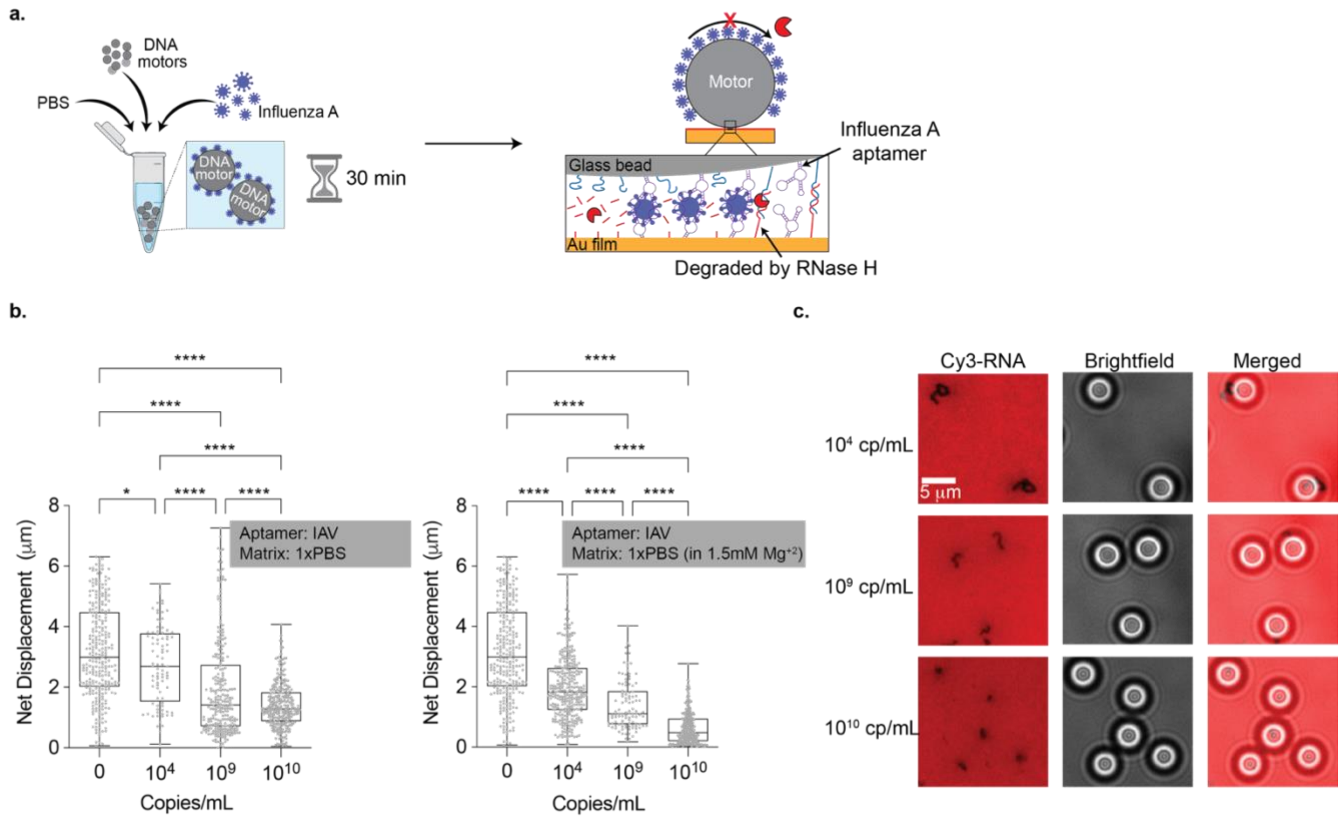


**Figure S12. DNA motors detect influenza A virus. a,** Schematic of sample generation for influenza A detection. DNA motors functionalized with 10% influenza A aptamer are incubated with influenza A virus in 1xPBS for 30 mins at room temperature. When added to the RNA chip decorated with 50% of the same aptamer, the motors stalled in the presence of RNAseH. **b,** Plot on the left shows the net displacement of >100 motors incubated with different concentrations of influenza A virus in 1xPBS. Plot on the right shows the net displacement of >100 motors incubated with different concentrations of influenza A virus in 1xPBS supplemented with 1.5mM $Mg^{+2}$. * and **** indicate $P$ =0.018 and $P$ < 0.0001, respectively. Experiments were performed in triplicate. **c,** Fluorescence and brightfield imaging of DNA motors with different concentrations of influenza A virus spiked in 1xPBS supplemented with 1.5 $Mg^{+2}$. RNA depletion tracks formed by the motors decrease as influenza A virus concentration increases.

**Figure S13. DNA motors tolerate breath condensate as a virus sensing medium.**

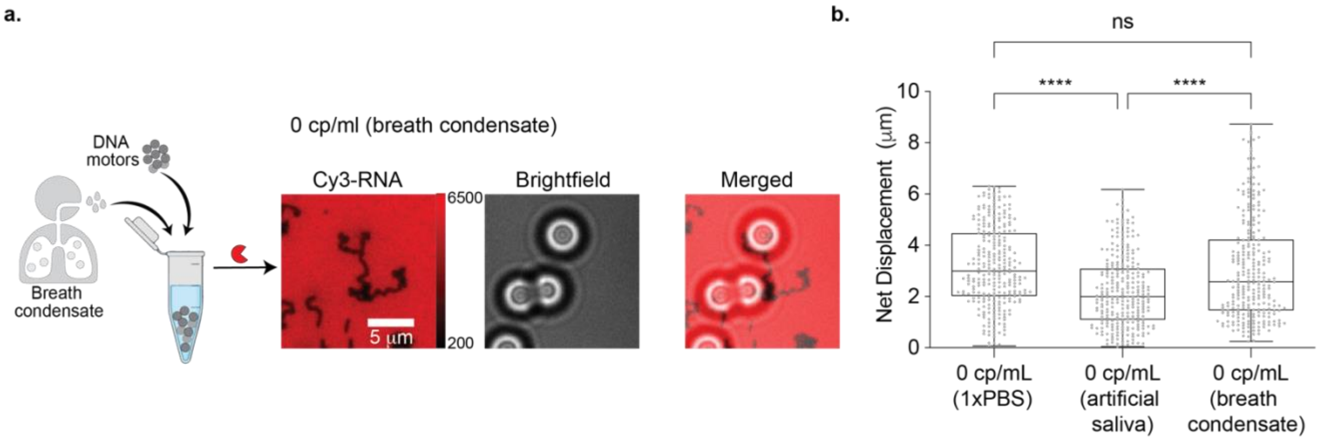


**Figure S13. DNA motors tolerate breath condensate as a virus sensing medium. a,** Schematic of sample generation in exhaled breath condensate. After collection, 1 μL of motors were mixed in the breath condensate sample. Fluorescence and brightfield imaging show that DNA motors diluted in breath condensate without virus display long depletion tracks on Rolosense chip with aptamer. The vertical color bar indicates the minimum and maximum values (a.u.) used to display the fluorescence of the Cy3-tagged RNA. We include this bar to note the integrity of the RNA monolayer. **b,** Plots of net displacement of ~300 motors with no virus diluted in 1xPBS, artificial saliva, and breath condensate. ns and **** indicate not statistically significant and $P < 0.0001$, respectively. Experiments were performed in triplicate.

**Figure S14. DNA-based motors stall in the presence of a single viral particle.**

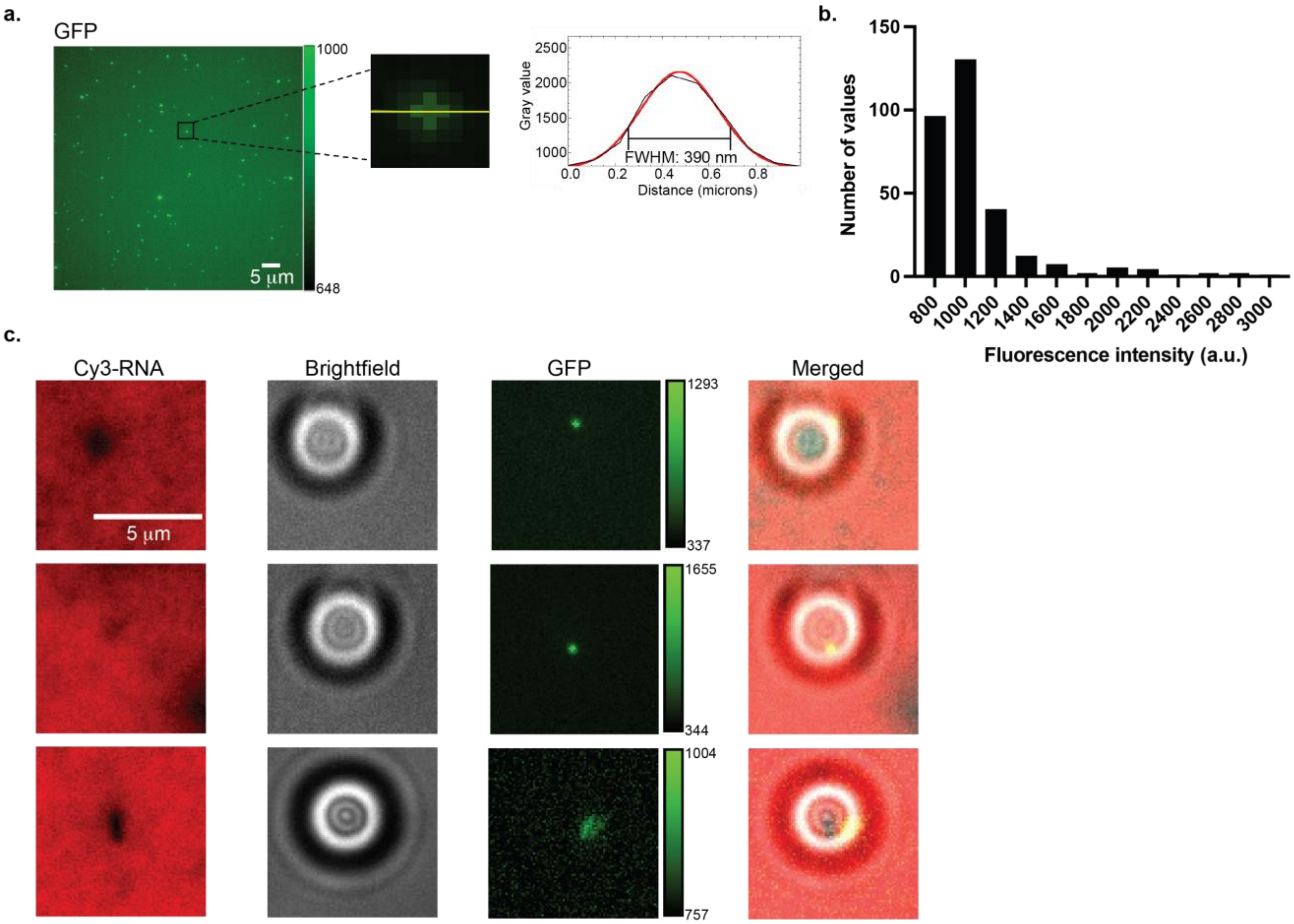


**Figure S14. DNA-based motors stall in the presence of a single viral particle. a,** Fluorescence imaging of 1000x diluted spike VLP particles (48 fM in 1xPBS) on the Rolosense chip. Line scan drawn on one of the particles indicate that the full width half maximum (FWHM) of a single particle is 390 nm. **b,** Distribution of the fluorescence intensity of 1000x diluted spike VLP particles in 1xPBS on Rolosense chip. Single VLP particles have a fluorescence intensity of 800-1200 a.u. **c,** Brightfield and fluorescence imaging of DNA motors incubated with 25 pM of GFP-labeled spike VLPs in 1xPBS. Samples with GFP-labeled spike VLPs show stalled motors and no depletion tracks along with a colocalized GFP signal with fluorescence intensity values indicating single particle. The vertical color bars indicate the minimum and maximum values (a.u.) used to display the fluorescence of the GFP-tagged VLPs. We include these bars to note the quantitative variations in fluorescence intensity of the bound VLPs.

**Figure S15. Rolosense chip can tolerate saliva in the presence of RNase inhibitors.**

**a.**

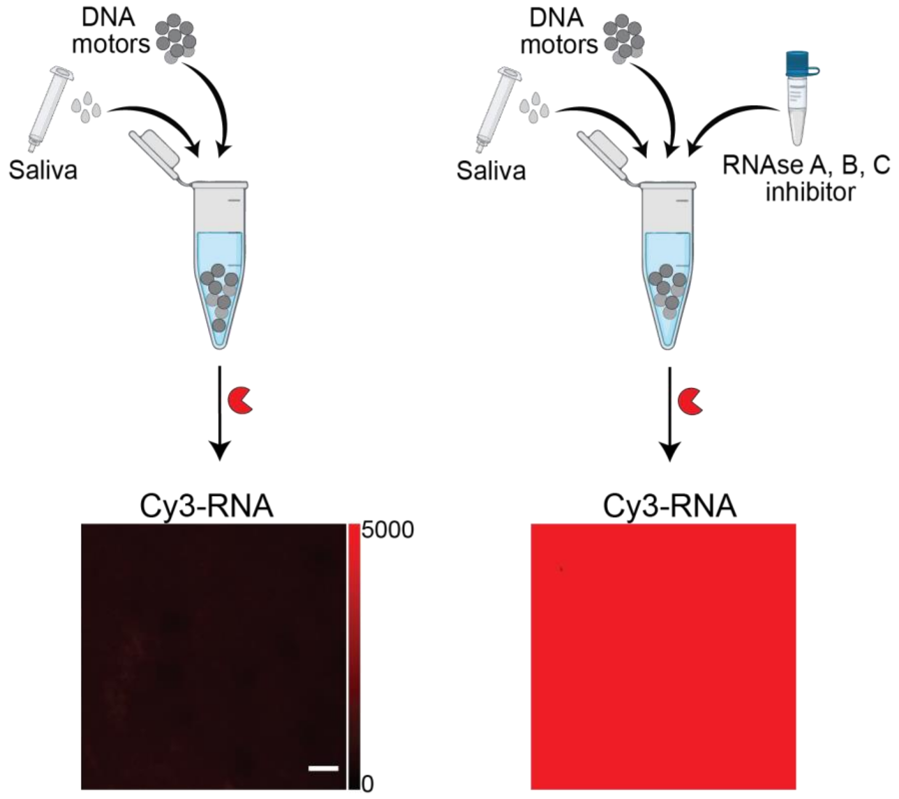


**Figure S15. Rolosense chip can tolerate saliva in the presence of RNase inhibitors.** **a,** DNA motors were incubated with saliva for 30 mins at room temperature and washed 3x in 1xPBS via centrifugation (15,000 rpm, 1 min). After wash the motors were added to Rolosense chip in the presence of RNaseH enzyme. Following 30 mins of RNaseH enzyme incubation, the Cy3 fluorescence intensity dropped significantly due to nuclease degradation. The Cy3 fluorescence intensity was recovered by incubating the motors for 30 mins at room temperature in saliva along with 20 units of RNase A, B, and C inhibitor. After the incubation, the sample was washed 3x in 1xPBS via centrifugation (15,000 rpm, 1 min) and added to the Rolosense chip along with RNaseH. The vertical color bar indicates the minimum and maximum values (a.u.) used to display the fluorescence of the Cy3-tagged RNA. We include this bar to note the integrity of the RNA monolayer following exposure to saliva. Scale bar is 10 μm.

**Supplementary Movies**

**Supplementary Movie S1:** Timelapse video of the brightfield channel superimposed with color-coded particle trajectories acquired at 5 s intervals for a duration of 30 mins. The video was acquired following RNaseH addition using a 20x 0.5 NA objective. SARS-CoV-2 and influenza A motors are not incubated with virus and both motors displayed long trajectories. The SARS-CoV-2 motors (6μm polystyrene) are distinguishable from the influenza A motors (5μm silica) on brightfield due to their darker outer rim. Scale bar is 10 μm.

**Supplementary Movie S2:** Timelapse video of the brightfield channel superimposed with color-coded particle trajectories acquired at 5 s intervals for a duration of 30 mins. The video was acquired following RNaseH addition using a 20x 0.5 NA objective. SARS-CoV-2 and influenza A motors are incubated with $10^{10}$ copies/mL of influenza A virus and only SARS-CoV-2 motors display long trajectories. The SARS-CoV-2 motors (6μm polystyrene) are distinguishable from the influenza A motors (5μm silica) on brightfield due to their darker outer rim. Scale bar is 10 μm.

**Supplementary Movie S3:** Timelapse video of the brightfield channel superimposed with color-coded particle trajectories acquired at 5 s intervals for a duration of 30 mins. The video was acquired following RNaseH addition using a 20x 0.5 NA objective. SARS-CoV-2 and influenza A motors are incubated with $10^7$ copies/mL of SARS-CoV-2 WA-1 and only influenza A motors display long trajectories. The SARS-CoV-2 motors (6μm polystyrene) are distinguishable from the influenza A motors (5μm silica) on brightfield due to their darker outer rim. Scale bar is 10 μm.

**Supplementary Movie S4:** Timelapse video of the brightfield channel superimposed with color-coded particle trajectories acquired at 5 s intervals for a duration of 30 mins. The video was acquired following RNaseH addition using a 20x 0.5 NA objective. SARS-CoV-2 and influenza A motors are incubated with $10^{10}$ copies/mL of influenza A virus and $10^7$ copies/mL of SARS-CoV-2 WA-1 and both motors remain stalled on the surface. The SARS-CoV-2 motors (6μm polystyrene) are distinguishable from the influenza A motors (5μm silica) on brightfield due to their darker outer rim. Scale bar is 10 μm.

**Supplementary Movie S5:** Representative timelapse smartphone video acquired at 5 s intervals for a duration of 15 mins using Cellscope. The video was acquired following RNaseH addition of DNA-based motors diluted in artificial saliva without virus present. Scale bar is 10 μm.

**Supplementary Movie S6:** Representative timelapse smartphone video acquired at 5 s intervals for a duration of 15 mins using Cellscope. The video was acquired following RNaseH addition of DNA-based motors incubated with $10^7$ copies/mL of SARS-CoV-2 B.1.617.2 spiked in artificial saliva. The motors were incubated with virus for 30 mins at room temperature and added to the Rolosense chip. Scale bar is 10 μm.
